# Ultra-high density imaging arrays for diffuse optical tomography of human brain improve resolution, signal-to-noise, and information decoding

**DOI:** 10.1101/2023.07.21.549920

**Authors:** Zachary E. Markow, Jason W. Trobaugh, Edward J. Richter, Kalyan Tripathy, Sean M. Rafferty, Alexandra M. Svoboda, Mariel L. Schroeder, Tracy M. Burns-Yocum, Karla M. Bergonzi, Mark. A. Chevillet, Emily M. Mugler, Adam T. Eggebrecht, Joseph P. Culver

## Abstract

Functional magnetic resonance imaging (fMRI) has dramatically advanced non-invasive human brain mapping and decoding. Functional near-infrared spectroscopy (fNIRS) and high-density diffuse optical tomography (HD-DOT) non-invasively measure blood oxygen fluctuations related to brain activity, like fMRI, at the brain surface, using more-lightweight equipment that circumvents ergonomic and logistical limitations of fMRI. HD-DOT grids have smaller inter-optode spacing (∼13 mm) than sparse fNIRS (∼30 mm) and therefore provide higher image quality, with spatial resolution ∼1/2 that of fMRI. Herein, simulations indicated reducing inter-optode spacing to 6.5 mm would further improve image quality and noise-resolution tradeoff, with diminishing returns below 6.5 mm. We then constructed an ultra-high-density DOT system (6.5-mm spacing) with 140 dB dynamic range that imaged stimulus-evoked activations with 30-50% higher spatial resolution and repeatable multi-focal activity with excellent agreement with participant-matched fMRI. Further, this system decoded visual stimulus position with 19-35% lower error than previous HD-DOT, throughout occipital cortex.

## 1. INTRODUCTION

Functional magnetic resonance imaging (fMRI) has enabled dramatic advances in cognitive neuroscience and human brain mapping.^1-6^ For example, in the last decade, fMRI has achieved remarkable success at decoding the identity and content of a wide range of stimuli experienced by human subjects.^7-10^ However, standard fMRI employs very large, non-portable equipment that cannot be used at the bedside, or with most patients with metal implants, is difficult in young children, and is not naturalistic. Functional near-infrared spectroscopy (fNIRS) images blood dynamics non-invasively, like fMRI, and has potential to address many fMRI limitations. While fNIRS traditionally used sparse imaging arrays and suffered from poor resolution and distorted point-spread functions, newer high-density diffuse optical tomography (HD-DOT) systems provide higher image quality and a superior surrogate to fMRI.^11-14^ Moving from sparse 30-mm spacing, with single-distance measurements, to high-density 13-mm spacing, with multiple-distance measurements, provides better brain specificity, improved resolution, and higher contrast-to-noise ratio. With HD-DOT, imaging is more comparable to fMRI in resting-state functional connectivity, retinotopy, and language tasks. However, it is unclear whether 13-mm spacing is optimal. Early simulation papers on DOT performance suggested resolution can linearly scale at 1/3-1/2x the depth into tissue,^15^ implying that at 15-mm depth (on gyri of the cortical surface), a resolution of 5-7 mm is possible. However, HD-DOT studies using 13-mm optode spacing have demonstrated only ∼13-16 mm resolution.^12^ Herein, we investigated the potential improvements of moving from 13-mm, to 10-mm, 6.5-mm, or 3.25-mm-spaced imaging arrays through simulations, experimental systems, and stimulus paradigms focusing first on imaging resolution and reliability, then on decoding visual cortex stimuli.

The challenges with increasing grid density include opto-electronic (e.g., dynamic range, crosstalk, detectivity with shorter dwell times while maintaining adequate frame rate), opto-mechanical (e.g., packing density vs. detector area, combing denser arrays through hair, scalp coupling), and computational (e.g., resolution of head models). While these challenges are widespread, they also present opportunities. For instance, while optode density scales with the square of the spacing, the number of measurements, or source-detector pairs (SD-pairs), scales with the fourth power of the spacing. Increasing the number of measurements often, ubiquitously, improves tomography systems.

We first evaluated imaging arrays with 13-mm, 10-mm, 6.5-mm and 3.25-mm spacing using high-resolution simulations. Simulated instruments were combined with physiologically based models of signal and noise to provide realistic performance predictions. Second, based on the simulation results, we designed, constructed, and tested a system with 6.5-mm spacing (ultra-high-density, or UHD). Relative to 13-mm, the 6.5-mm system required several advances in electro-optics, including expanding the dynamic range by more than 30x to 3x10^8^, diminishing crosstalk by an equivalent amount, and retaining the same effective noise floor while reducing dwell time by 10x. We then evaluated the fidelity of the 6.5-mm system with retinotopic mapping paradigms and evaluated these images against participant-matched fMRI. Finally, we evaluated the information-transfer/bit-rate perspective by analyzing how grid density affected the accuracy of decoding stimulus position from brain activity in visual cortex. Collectively, these studies demonstrate that 6.5-mm arrays significantly improve resolution laterally and in depth, increase signal-to-noise-ratio, and enable higher-performance decoding of optically measured brain function, setting new benchmarks for non-invasive optical neuroimaging system designs.

## 2. RESULTS

### 2.1 Simulation Studies

To assess potential improvement in image quality, we reconstructed images from simulated DOT measurements at grid spacings of 13 mm, 10 mm, 6.5 mm, and 3.25 mm (**Fig. 1A-D**). Simulated measurements were generated using a forward model, *y* = *Ax* +*n*, relating the source-detector differential measurements (*y*) to the product of the light model Jacobian matrix (*A*) and a hypothetical absorption image (*x*), to which we added light intensity-scaled noise (*n*). The Jacobian was created using a finite-element calculation from the diffusion approximation. To evaluate the position-dependent point-spread functions (PSFs), 105,175 single-voxel absorption targets were imaged throughout the field-of-view. To match the signal-to-noise ratio in simulations to in-vivo measures, we scaled the targets’ amplitude to match typically measured variance in *y* (see **Supplementary Information**),^13^ which captured physiology-based and instrument noise. Images were reconstructed using standard linear inverse methods with 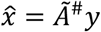. Here, 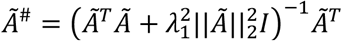, a spatially-variant-Tikhonov-regularized pseudo-inverse of *A* with regularization parameter *λ*_1_.^13^ The spatially-variant regularization provides a more-even response versus depth.^13,16^ PSF images were evaluated for their width, position, and repeatability (**Fig. 1-2**).

**Figure 1:**
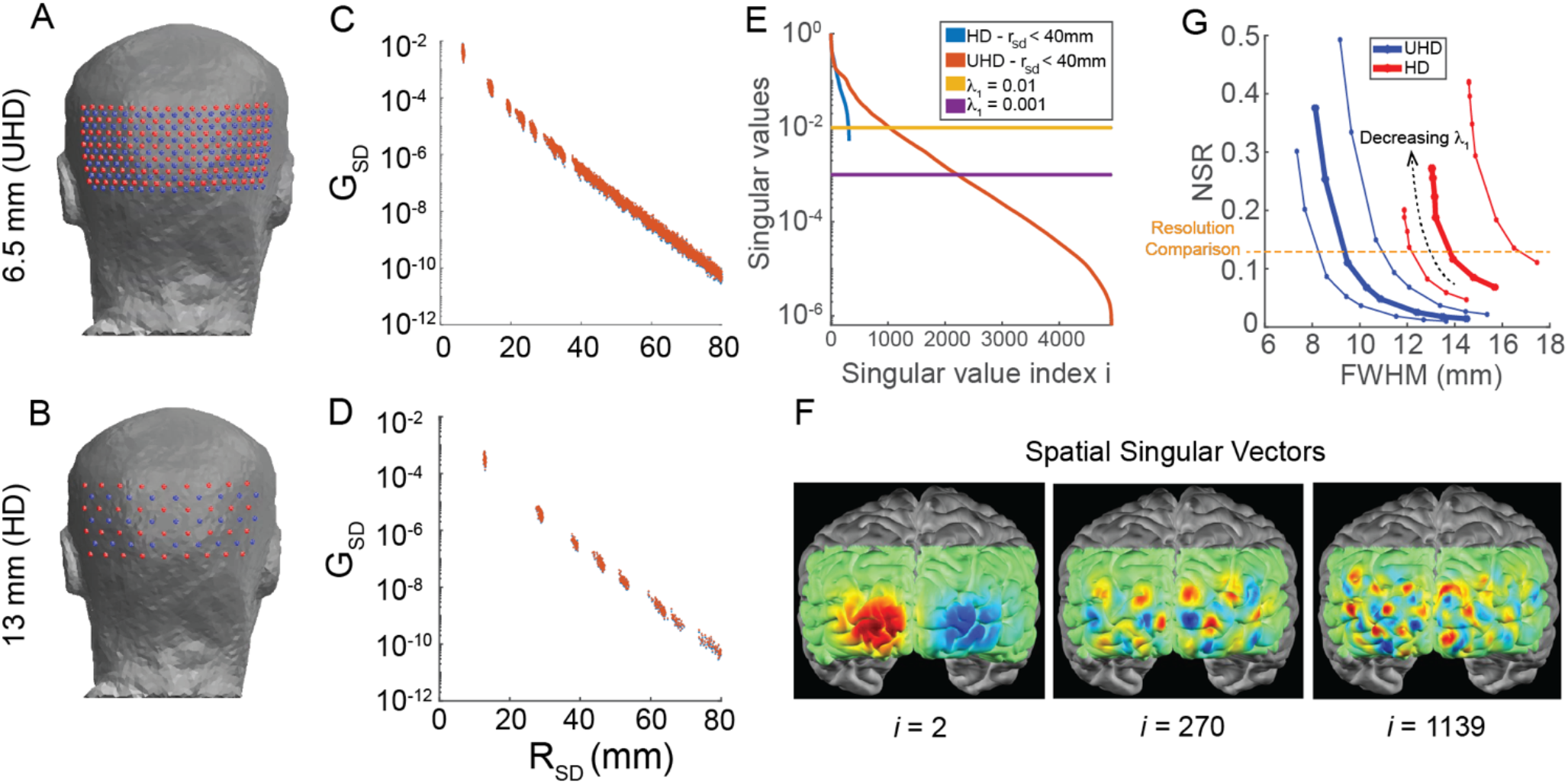
Simulated DOT image quality from grids with different nearest-neighbor spacing (6.5 mm and 13 mm). **(A-B)** Grids on head model. **(C-D)** Simulated baseline light level vs. source-detector distance follows the expected log-linear decay. **(E)** Singular values of each grid’s Jacobian matrix. The 6.5-mm grid contains more singular values, and these values decay far less quickly, which should give the 6.5-mm grid a better noise-resolution tradeoff as *λ*_1_ varies. **(F)** *λ*_1_ creates a tradeoff between image resolution and noise amplitude because singular vectors with higher singular values contain lower-spatial-frequency patterns. **(G)** Simulated noise-to-signal ratio (NSR) vs. FWHM of reconstructed point target images for multiple levels of regularization (*λ*_1_ values 0.001, 0.0033, 0.005, 0.0066, 0.01, 0.033, 0.066, 0.1). As *λ*_1_ decreases, resolution improves (FWHM decreases) at the expense of increasing NSR (dotted black trend line). For a given NSR (e.g., orange dashed line), the 6.5-mm grid has superior resolution.

To establish whether a system has better general imaging performance, *λ*_1_ must be swept to evaluate resolution at comparable noise levels (**Fig. 1E-G**). First, we examined the singular values of each system’s Jacobian, excluding channels with source-detector distance > 40 mm. The 13-mm (HD) grid runs out of nonzero singular values when *λ*_1_∼0.005-0.01 (**Fig. 1E**) indicating that the resolution limit will be reached at *λ*_1_∼0.005-0.01 (**Fig. 1G**). In contrast, singular values for the 6.5-mm (UHD) grid decay more slowly and have a much-lower limit (**Fig. 1E**), suggesting that the resolution will continue improving as *λ*_1_ decreases. Evaluating PSFs across varying noise-to-signal ratio (NSR) levels (**Fig. 1G**), the full width at half maximum (FWHM) was 30-50% smaller for the UHD grid than HD, establishing that in simulation the UHD grid has a superior noise-resolution tradeoff, due to the UHD grid’s ability to incorporate singular vectors with higher spatial frequencies (**Fig. 1F**).

These simulations reveal that increasing the grid density from 13-mm to 6.5-mm spacing should improve image quality (**Figs. 1G, 2**). PSFs from the 6.5-mm (UHD) grid had FWHM 30-50% smaller than from the 13-mm grid at most depths and with better regularity, indicating higher and more consistent spatial resolution from the 6.5-mm grid (**Fig. 2A-E**). Also, PSF centroids from the 6.5-mm grid had smaller localization error (**Fig. 2A,E**). PSFs’ signal-to-noise ratio (SNR) also increased with grid density. In summary, these simulations predict that increasing grid density from 13-mm to 6.5-mm spacing should produce improvements of 5-7 mm in spatial resolution, 2-4 mm in localization error, 1.5-2.0x in effective spatial resolution, and 1.4-2.0x in SNR when imaging 8-20 mm deep in the head (**Fig. 2G**). The simulations predict more-modest resolution improvements from increasing grid density to 3.25-mm spacing.

**Figure 2:**
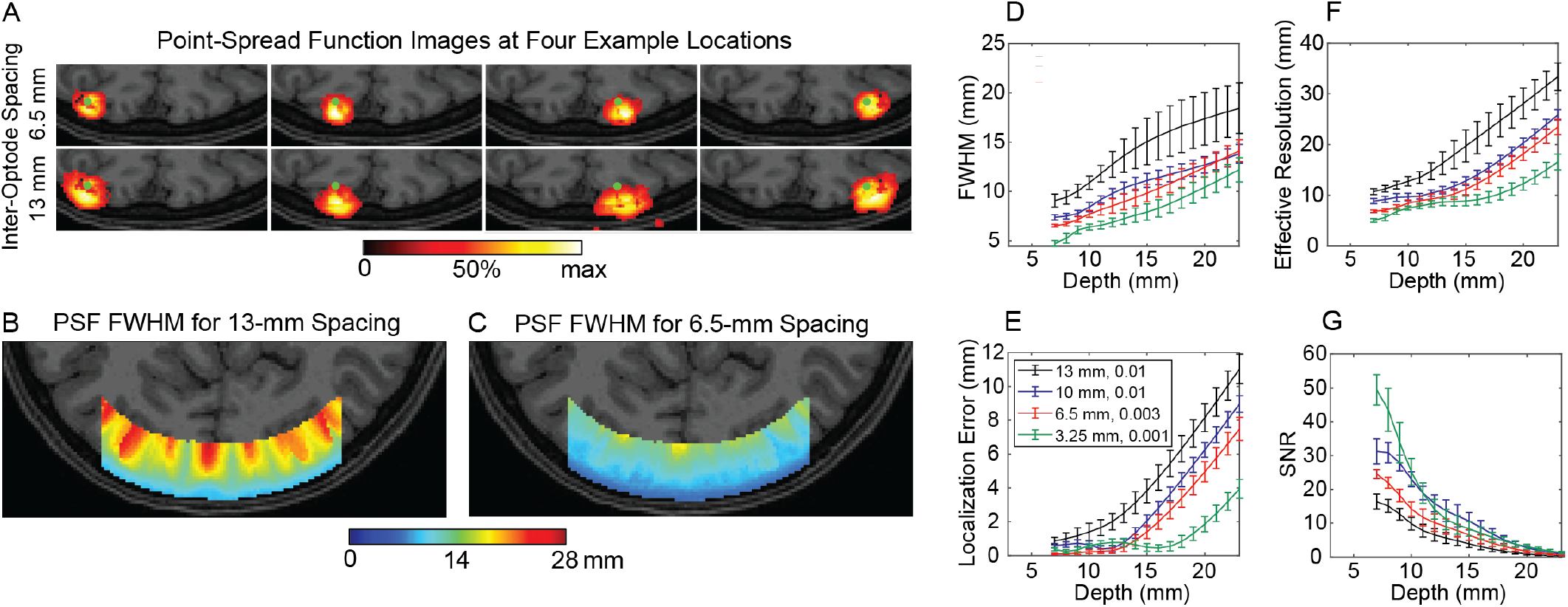
Simulated DOT image quality and its depth dependence for grids with different nearest-neighbor spacing. **(A)** Point-spread functions (PSFs) of single-voxel (point) targets in the head, using simulated measurements. PSFs for the 6.5-mm grid have smaller spatial extent, are less noisy, and are closer to their point targets. **(B)** PSF FWHM throughout a region of interest for the 13-mm grid. **(C)** PSF FWHM throughout a region of interest for the 6.5-mm grid. **(D-G)** Metrics of the PSF, including FWHM **(D)**, localization error **(E)**, effective resolution **(F)**, and SNR **(G)** vs. depth of point target, for four spacings and *λ*_1_ values. Error bars denote standard error of the mean over locations. These metrics of image quality improve as inter-optode spacing decreases at all depths of interest. However, the improvements from decreasing 6.5-mm to 3.25-mm spacing were far smaller than from decreasing 13-mm to 6.5-mm spacing.

### 2.2 UHD-DOT Instrument

Motivated by the simulations, we constructed a 6.5-mm UHD-DOT system with 126 sources, 126 detectors, and 2 wavelengths (**Fig. 3a-b**). The two main challenges for a 6.5-mm vs.13-mm system were to: 1) increase dynamic range by 30x to accommodate higher light levels in 6.5-mm-spaced source-detector pairs, and 2) triple light levels to compensate for the lower dwell time necessary for 91 instead of 16 time-encode steps; HD-DOT systems use spatial, temporal, and frequency encoding to separate detector signals into source-detector signals. To triple light levels, previous LED illumination was replaced by laser diodes. To increase the dynamic range in light levels, a two-pass encoding scheme was employed (**Fig. 3b-c**). The first (“bright”) pass used 50% duty cycle illumination for each source time step and was optimized for farther-away detectors. The second (“dim”) pass used 1% duty cycle and was optimized to avoid saturating nearby detector circuits. The dim measurements were scaled to equivalent of 50% duty cycle. The rest of the system leverages the design of previous HD-DOT systems^13,17^ (see **Supplementary Information** for full details). The 6.5-mm system yielded 4,917 source-detector-pair measurement channels (for each wavelength) usable for image reconstruction (i.e., channels with source-detector separation ≤ 40 mm), a ∼20x greater channel count than 13-mm grids covering the same region.

**Figure 3:**
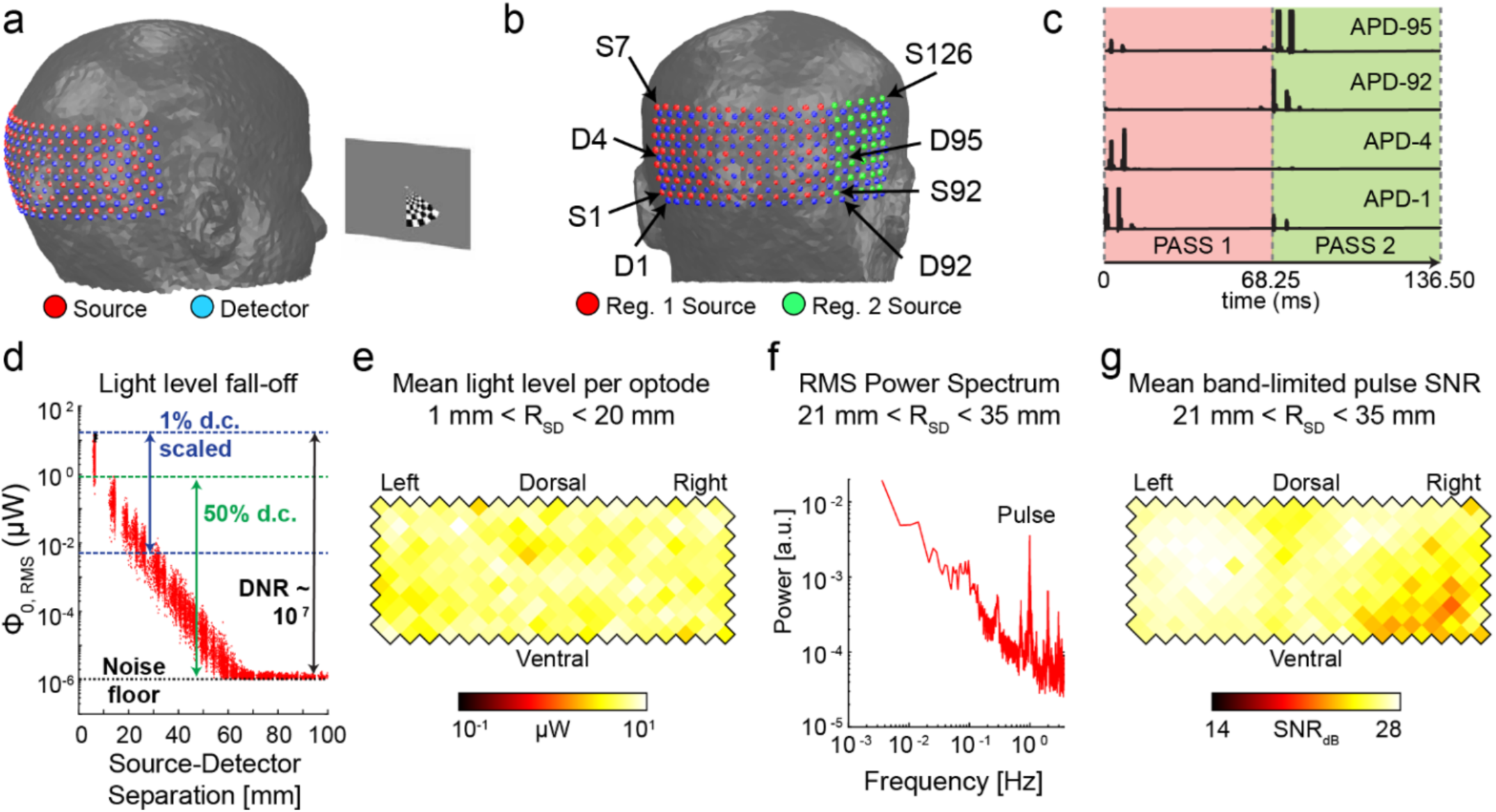
Ultra-High-Density Diffuse Optical Tomography system with 6.5-mm inter-optode spacing. **(a)** Diagram of human brain imaging experiment: human subject views checkerboard stimuli while the grid of sources and detectors is placed on the back of the head. **(b)** Grid with the two regions used for two-pass encoding. **(c)** Raw signals from different detectors in two-pass encoding. **(d)** Light (830 nm) fall-off curve from human subject decays with expected log-linear fashion with source-detector pair distance and indicates a dynamic range of ∼10^7^ between the saturation ceiling and noise floor. **(e)** Mean light level in superficially-sampling measurements involving each optode, displayed on a flattened grid, indicating strong, even optode-to-scalp coupling. **(f)** Cardiac pulse peak at ∼1 Hz in the root-mean-square (RMS) averaged power spectrum (source-detector pair distances 21-35 mm). **(g)** Cardiac pulse power signal-to-noise ratio (SNR) exceeds 20 dB for each optode.

The UHD-DOT system achieved a dynamic range (DNR) of ∼3x10^8^, spatially consistent optode-to-scalp coupling, and high cardiac pulse SNR (∼20 dB) (**Fig. 3d-g**). The light fall-off curve of the mean light level vs. SD-pair distance follows the expected log-linear form (**Fig. 3d**) and reveals the noise floor, dynamic range, and crosstalk performance. The noise floor for single time step is 1.0 pW at 910 Hz, or noise-equivalent power 10 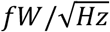 at 1 Hz, matching the avalanche photodiode specification. At short source-detector distances, detection modules are at their saturation ceiling. The ratio of saturation ceiling to noise floor defines the DNR, which is 10^7^ at 910 Hz, or 3x10^8^ at 1 Hz. Cap fit is optimized by assessing the mean light levels for each optode from source-detector pairs with distance < 20 mm, which are minimally sensitive to tissues deeper than the skull (**Fig. 3e**). Cardiac pulse, generally the strongest physiological signal for head-mounted near-infrared measurements, provides a useful, fast measure of data quality (**Fig. 3f**). Cardiac pulse SNR (power at pulse frequency vs. nearby frequencies) is strong across the entire cap (**Fig. 3f-g**).

### 2.3 Noise-Resolution Tradeoff Evaluation with Head Phantom and In-Vivo Data

To assess noise vs. resolution in our UHD-DOT (6.5-mm spacing) instrument, we compared images using the full 6.5-mm-spaced grid to a subset of those channels at 13-mm spacing (“HD subset”). We evaluated phantom and in-vivo data using three approaches (**Figs. 4-5**). In the first approach, we imaged a head phantom to evaluate the impact of electro-optic noise without physiologic noise. As in simulation studies, we compared image noise to resolution while varying λ_1_. Image noise was estimated from the phantom data, while resolution and signal amplitude were estimated from the FWHM of simulated PSFs. FWHM remained 30-50% lower for the full grid than the HD-subset grid at each *λ*_1_ value (**Fig. 4B**) and for comparable NSR values (**Fig. 4D**). Resolution and NSR also behaved as expected; for each grid, PSF FWHM increased and NSR decreased as *λ*_1_ increased (**Fig. 4B-C**), with minimal change in HD-DOT results for *λ*_1_<0.01.

**Figure 4:**
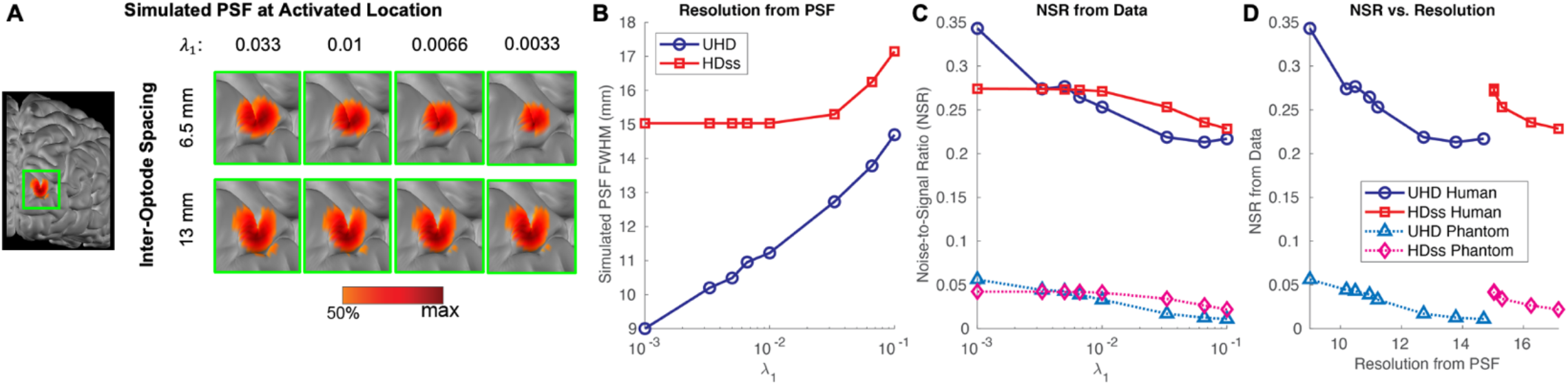
Resolution (simulation) vs. noise (measured). (**A**) Simulated PSF on the brain surface for different levels of regularization for the full UHD grid (6.5-mm spacing) and its HD-subset (13-mm spacing) for a representative visual cortex voxel. (**B-C**) As *λ*_1_ increases, the PSF FWHM increases and NSR decreases, as expected, creating the noise-resolution tradeoff in panel **D**. The NSR was calculated within the FWHM-activated region of a human subject viewing a rotating checkerboard wedge. For the static head phantom, data measured on the head phantom were reconstructed into images using a generic human head model. Importantly, the system noise is ∼5x lower than in-vivo noise highlighting how low the noise floor is with this system. (**D**) Experimental NSR vs. simulation resolution tradeoff. At a given NSR, the resolution of the UHD grid is superior to the HD-subset grid for both the phantom and human subject cases.

**Figure 5:**
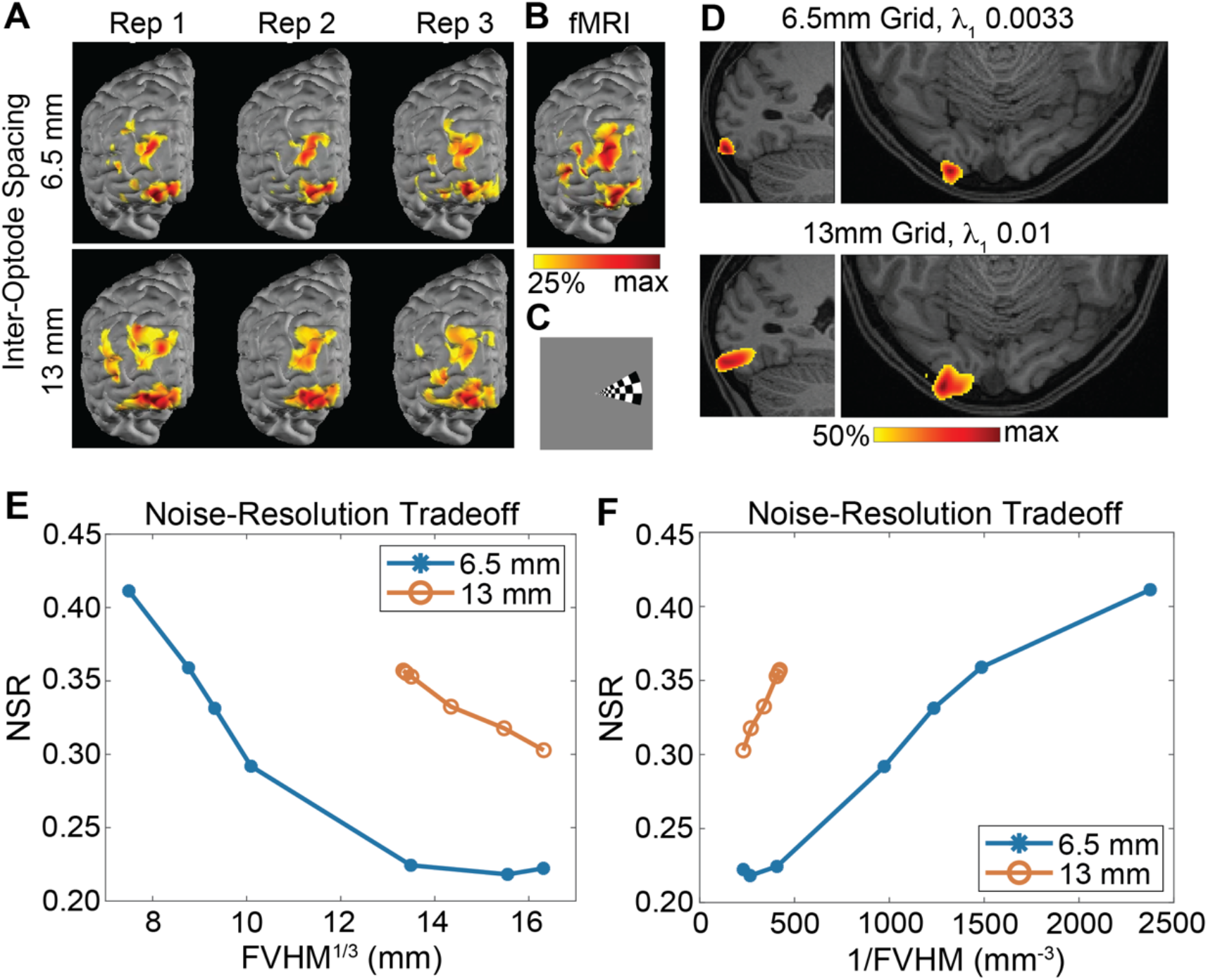
In-vivo imaging validation. **(A-B)** Multi-cluster oxyhemoglobin activation images (*t*-maps) evoked by the counterclockwise-revolving checkerboard wedge stimuli, as imaged by the UHD and HD-subset grids and fMRI. Each column of panel **A** was formed by block-averaging the images from 1/3 of the repetitions of one wedge position (panel **C**). Across these three repetition sets, the images were repeatable and resembled fMRI, and the activation clusters were wider in the 13-mm array images than 6.5-mm array images. The color scale’s upper bound is the maximum value in the activation image from the UHD grid over all stimulus repetitions. **(D)** Volume-slice views of the response to one wedge position show a smaller activation for 6.5-mm vs 13-mm imaging array, that is also better localized within brain tissue. The color scale’s upper bound is the maximum value within each transverse slice. **(E-F)** Noise-resolution tradeoff, illustrated by curves of peak NSR vs. FVHM^1/3^ and peak NSR vs. 1/FVHM of the stimulus-evoked response using various *λ*_1_. At any given NSR, the activation FVHM^1/3^ remained 30-50% smaller and 1/FVHM remained 3-5x larger for the UHD grid than HD-subset, indicating UHD’s superior noise-resolution tradeoff.

In a second approach, we evaluated the effect of grid density with both electro-optic and physiologic noise. Noise was estimated from human images, while signal and resolution were again taken from simulated PSFs. The noise-resolution tradeoff for the human subject is more similar for the two grids than for the head phantom (**Fig. 4D**), but as *λ*_1_ decreases, the UHD grid provides higher resolution than its HD subset without a sharp NSR increase until *λ*_1_<0.0033 (**Fig. 4D**). In-vivo NSR exceeded phantom NSR by ∼5x. This is expected given that physiological noise dominates typical system noise by roughly a factor of five.

In the third approach, we evaluated both noise and resolution using in-vivo functional imaging data. A human participant was scanned while viewing flickering, revolving checkerboard wedges and flickering, expanding and contracting checkerboard rings that evoked brain activity peaks in visual cortex (**Fig. 5**). Resolution was quantified by the cube root of the volume (FVHM^1/3^) of the contiguous region containing the peak and having intensity ≥50% that peak inside an activation SNR image. The activation SNR image was formed by each voxel’s average signal value divided by that value’s standard deviation over stimulus repetitions (**Fig. 5D**). As *λ*_1_ was swept, FVHM^1/3^ remained 30-50% lower and 1/FVHM (roughly proportional to space-bandwidth product^18^) remained 3-5x higher for the UHD-DOT grid than HD subset at each NSR value (**Fig. 5E**).

### 2.4 Visual Cortex Response Repeatability, Resolution, and Comparison to fMRI

Due to repetition of the retinotopic visual map for each of the hierarchical functional areas in primary visual cortex, many in-vivo retinotopic responses are more complex than single-site PSFs, wherein the stimulus-evoked activation appears to split into multiple distinct clusters^19^. This provides an opportunity for evaluating resolution because a higher-resolution imaging system should see this splitting, whereas a lower-resolution system will blur the clusters together.

To validate these complex DOT images, we assessed their repeatability across different subsets of the data; we split the data into three equal-duration epochs and computed block-averaged oxyhemoglobin activation *t*-maps for each epoch (average signal divided by its standard error in each voxel) (**Fig. 5A**). The UHD-DOT images consistently show finer detail and smaller activation regions than their HD-subset counterparts. The consistency of these maps across data subsets also assuages the potential concern that these splitting effects might be due to noise.

To obtain a “ground-truth”/gold-standard measure of the evoked activity, we acquired fMRI data while subjects viewed rotating checkerboard wedges. At some timepoints, the fMRI images display the anticipated splitting into multiple clusters. At these timepoints, this splitting is also visible in the UHD and HD-subset images, though the clusters are wider/blurrier for HD-subset (**Fig. 5A-B**).

As further validation, we estimated retinotopic maps from our DOT images. Because our stimuli were periodic in time, stimulus phase corresponds to stimulus position, and the phase of a voxel’s activation at the stimulus repetition frequency indicates which stimulus position most strongly activates that voxel’s tissue. Images of the polar angle and eccentricity of these positions were derived from each voxel’s response to the wedges and rings, respectively, as in previous work (**Fig. 6A,C**).^12,19^ These maps followed the expected retinotopic organization of visual cortex areas.^19,20^

**Figure 6:**
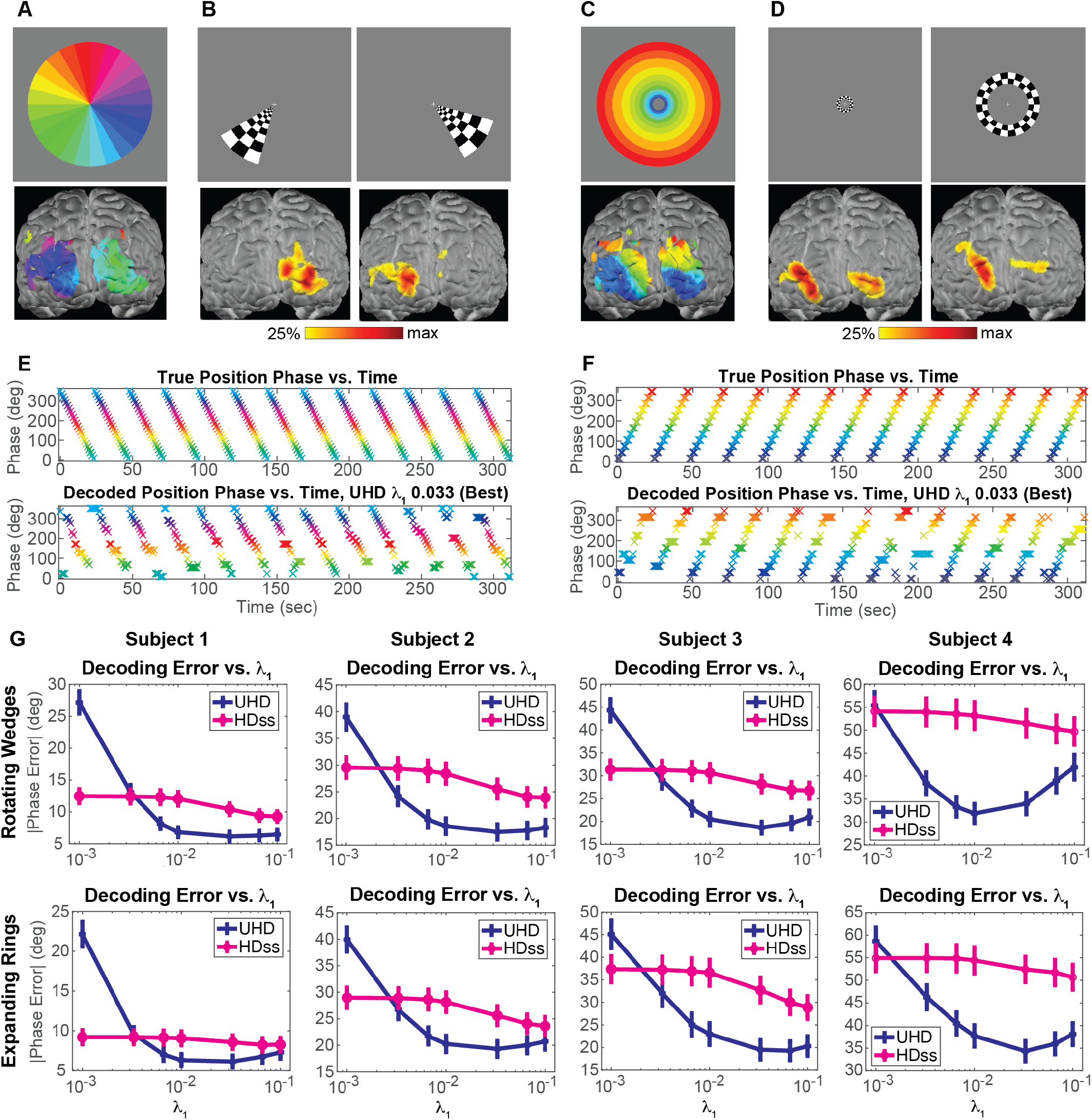
UHD-DOT outperforms HD-DOT in stimulus position decoding. **(A)** Phase-encoded retinotopic map of wedge stimulus position to which each voxel was most responsive. **(B)** Oxyhemoglobin activity evoked by two wedge positions, block-averaged over the first 26 repetitions of the counterclockwise-revolving wedge. **(C-D)** Analogous to panels A-B but for ring stimuli. Brain images in panels B and D were four of the decoder training templates. **(E)** True and UHD-decoded stimulus phase vs. time in one run with 13 counterclockwise wedge revolutions, for Subject 3 and the λ_1_ value that yielded lowest decoding error. **(F)** Analogous to panel E but for expanding rings. **(G)** Overall mean absolute error in decoded phase. Error bars denote standard error of the mean. For *λ*_1_≥0.01, the UHD grid produced significantly lower decoding error than HD-subset (HDss) in each subject. For *λ*_1_ values that yielded best decoding, decoding error was 19-35% lower for UHD than HDss.

### 2.5 Neural Decoding of Stimulus Position

A challenge with evaluating resolution in vivo is that the evoked activations are more spatially complex than simple points (**Fig. 5A-B**). Neural decoding tasks are also sensitive to the spatiotemporal information content (space-bandwidth product) and SNR of an imaging system and so provide a useful data-driven metric of image quality. Therefore, we evaluated the performance of the UHD and HD-subset grids at discriminating the position/phase of checkerboard wedges and rings using a simple brain activity decoding test^21^.

Images from the 6.5-mm grid yielded significantly higher stimulus position decoding accuracy than the 13-mm subset (**Fig. 6**). Specifically, mean absolute error in stimulus phase decoded by the UHD grid was 19-35% lower than HD-subset for 0.0066 ≤ *λ*_1_ ≤ 0.1, the regime of best decoding performance. These differences were also statistically significant (*p*<10^-4^, one-sided paired/matched-samples permutation test). Also, the *λ*_1_ values that maximized decoding accuracy differed from those which produced the highest-resolution images. This is neither surprising nor problematic. Rather, for these simple 24-way and 12-way decoding tasks, higher SNR (from larger *λ*_1_) may be more important than high spatial resolution for decoding accuracy.

## 3. DISCUSSION

The impact of DOT in neuroscience, particularly as a surrogate for fMRI, will be determined by how closely DOT image quality approaches that of fMRI. Herein we evaluated the potential improvements in DOT image quality through increasing the imaging grid density beyond 13-mm to 10-mm, 6.5-mm and 3.25-mm spacing. Simulations with noise predicted significant improvements for 10-mm and 6.5-mm, with diminishing returns seen beyond 6.5-mm (**Fig. 2**). After constructing a 6.5-mm system, with concomitant hardware improvements needed, we found that in-vivo resolution and noise performance matched trends from simulations, and resolution improvements were validated using participant-matched fMRI and repeatability analysis. Further, beyond raw image quality improvements, the UHD system significantly outperformed its HD subset in decoding retinotopic-mapping stimulus positions.

Motivated by prior studies that demonstrated image quality improvement from HD grids over sparse grids,^11,12^ we compared UHD and HD grid performance in simulation. Simulations predicted better noise-resolution tradeoff and localization error by moving from 13-mm to 6.5-mm inter-optode spacing. The 13-mm-spaced grid ran out of nonzero singular values as λ_1_ decreased beyond 0.0066, not far below the typical 0.01 value that has previously worked well with a 13-mm-spaced grid^12,13^. For equivalent noise levels, 6.5-mm vs. 13-mm grids showed better resolution by ∼4x in PSF volume and 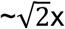 in linear resolution (FWHM) (**Fig. 1**). These findings aligned with and confirm previous simulations predicting superior performance from decreasing inter-optode spacing from 30 mm to 13 mm ^11^ and a previous in-vivo finger stimulation study that produced smaller, less-overlapping activation images when inter-optode spacing decreased from 15 mm to 7.5 mm.^22^

Though challenging due to requirements for opto-electronic sensitivity and noise, dynamic range, and ergonomics, we established the feasibility of building a 126 source x 126 detector UHD-DOT system covering a 120-mm x 55-mm area over visual cortex. Basic data quality metrics including light level decay vs. source-detector distance and cardiac pulse SNR established proof of principle for this design to measure neural hemodynamics with significant improvements over previous state-of-the-art systems. This system yielded 4917 measurement channels usable for image reconstruction per wavelength, ∼20x the number of channels provided by HD-DOT grids covering the same area. To extend the dynamic range, two-pass illumination enabled separate optimization of signal-to-noise ratio for short vs. long-distance source-detector pairs and provided a dynamic range ∼10^7^ (140 dB) (**Fig. 3**). With two-pass encoding and previously developed illumination multiplexing schemes the imaging frame rate (7.33 Hz) remained more than sufficient to track hemodynamic responses.^13,14,17,23^

In-vivo performance followed the trends predicted by simulations. Evaluations of retinotopic-mapping stimuli, showed image quality improvements with UHD-DOT through performance on resolution-noise tradeoff curves, comparison to fMRI images, repeatability analysis, and stimulus decoding. UHD-DOT improved linear resolution by 30-50% at a given noise level, and yielded sharper images that tended not to blur nearby activation clusters together as much. These features were repeatable over multiple subsets/splits of the dataset (**Fig. 5**). UHD-DOT images also produced 19-35% higher accuracy in decoding the position (phase) of the periodic stimuli (**Fig. 6**). Previous work has shown feasibility of reconstructing somatosensory responses with 7.5-mm inter-optode spacing, but the response images were not evaluated for noise-resolution tradeoff and were confined to a smaller, single-hemisphere field of view^22^. Our work contributes a more-comprehensive evaluation of these factors, a larger field of view, and more discussion of the theoretical support for the higher performance of UHD-DOT. Together, these studies demonstrate the promise of UHD-DOT for achieving higher image quality and information content for mapping and decoding.

Portability and wearability are major strengths of commercial optical neuroimaging devices with much-sparser grids. For wearability of fiber-based fNIRS/DOT, including our UHD-DOT system, the number of fibers determines the effective weight. In this context, 6.5-mm-spaced systems are 4x heavier than 13-mm-spaced systems. However, emerging fiberless and/or wireless DOT systems do not suffer from the same weight/density penalty^24-27^. Recent fiberless systems have replicated some capabilities of fiber-based systems, including retinotopic mapping^25^ and adult and infant resting-state network mapping^28,29^. This study highlights the importance of maintaining high optode grid density in these wearable systems.

In conclusion, herein we quantitatively evaluated the potential performance boosts from increasing the density of DOT imaging grids, showing significant increases image quality and information content for mapping and decoding tasks. These results suggest that UHD-DOT will yield strong decoding performance with more-complex, naturalistic visual stimuli including movies^9,30^ and point to the future performance of widely available wearable DOT systems.

## METHODS

### M.1. Simulation Studies

To examine the effect of optode grid density on imaging performance, simulations were performed with four DOT optode grid densities of nearest-neighbor spacing 13 mm, 10 mm, 6.5 mm, and 3.25 mm. To assess key measures of image quality, we simulated images of single-voxel activations at 105,175 locations within the head. The resulting point spread functions were quantitatively assessed for resolution, positional accuracy, and signal to noise, and evaluated as a function of depth.

Simulating these high grid densities presented computational challenges. For a given field of view or area covered by the optode grid, the numbers of sources *N*_*src*_ and detectors *N*_*det*_ each scale with the inverse square of the nearest-neighbor spacing *D*_*NN*_. The number of source-detector pair measurements, scales with the product *N*_*src*_*N*_*det*_ . Thus, the number of measurements scales with 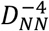, or equivalently with the square of the optode grid density. This causes the number of measurements to grow rapidly as the nearest-neighbor spacing is reduced. Also, accurately evaluating images from grids with smaller inter-optode spacing requires finer spatial sampling of the tissue (i.e., smaller voxels). For a given volumetric field of view and isotropic voxels, the number of voxels scales with the inverse cube of the voxel edge length.

To manage this computational explosion while retaining adequately small voxels (cubes with edge length 1 mm), we restricted our optode grid size (coverage area) and field of view. For all nearest-neighbor spacings considered, the optode grid covered a 120 mm x 55 mm region over the back of the head. Head tissue farther than 40 mm from the optode grid was ignored in all modeling because at that distance, the light intensity from a source would have decayed to 10^-5^ times the intensity at the source or less, for the wavelengths of interest for imaging (685-830 nm). Measurements whose source-detector distances exceeded 40 mm typically have insufficient light levels resulting in poor signal-to-noise and so were excluded ^13^. Simulated image quality analyses were then further restricted to a slab subregion near the center of the optode grid. This subregion covered a left-right length of 100 mm, a superior-inferior height of 35 mm, and depths 5-25 mm into the head, which excluded scalp tissue and deep brain tissue. Image quality in DOT varies strongly with depth but is largely shift-invariant in lateral directions ^15^, and therefore the findings of these simulation studies are expected to generalize to optode grids with similar densities but larger areas of coverage. To test image quality, we placed simulated point activations at each voxel in the subregion defined above.

#### M.1.1. Imaging System Forward Model

Simulations employed an established linear forward model for continuous-wave DOT systems ^13,17,31,32^. Briefly, this model relates changes in the optical absorption coefficient 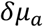 at each tissue location (volumetric voxel or mesh node) to the log-ratio fluctuation in measured light levels Φ about their baseline temporal mean Φ_0_ for each source-detector pair. These quantities are related by the following equation, where *v* is the speed of light in the tissue, *D* is the optical diffusion coefficient, and *G*(*r’, r, λ*) is the light fluence induced at position *r* by a point emitter at position *r’* with light wavelength *λ*. Formally, *G*(*r’, r, λ*) is the Green’s function solution to the steady-state optical diffusion equation.

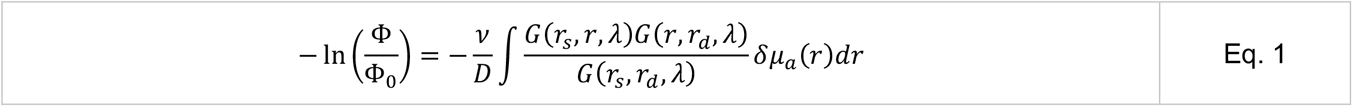

Approximating the integral for each measurement channel *i* as a discrete sum over voxels *j* and making the following definitions (Eq. 2), we arrive at a linear forward model of the imaging system in the absence of noise (Eq. 3):

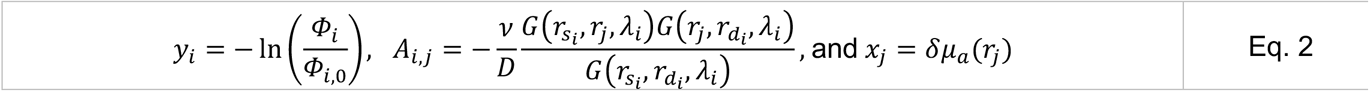

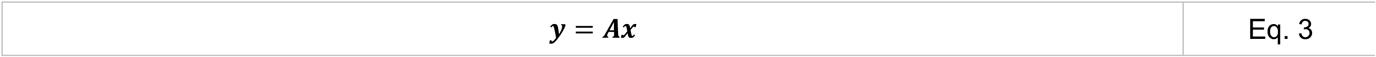

Each row of the Jacobian matrix **A** corresponds to the measurement *y*_*i*_ made by a source-detector pair at wavelength *λ*_*i*_ and encodes the spatial sensitivity profile of that measurement to absorption changes in the tissue. To generate **A** for a given grid of sources and detectors, the grid was positioned on the back of a segmented human head mesh. The head mesh was constructed from a T1-weighted MRI image using the software packages FreeSurfer ^33^ and NeuroDOT ^34,35^, as previously described ^13^. The Green’s functions were then computed using the software package NIRFAST, which employs a finite-element model solver ^36-38^.

To incorporate the important presence of noise **n** in real measurements, we employed the following forward model of the imaging system that relates the measurements **y** to the image **x** through the light model sensitivity matrix **A** :

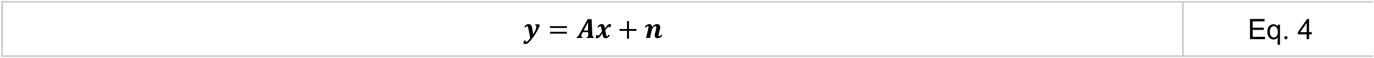

The noise **n** was simulated from a simplified model fitted to data from a previous DOT system (Eggebrecht et al. 2014). In this noise model, **n** follows a multivariate Gaussian distribution where each measurement channel’s variance depends on the source-detector separation and the covariance between channels is 0. This empirical noise model captures variance related to both physiology and shot noise. Further details and fitting procedures for this noise model are described in **Supplementary Information**.

#### M.1.2. Image Reconstruction (Inverse Problem)

Image reconstruction employed a previously established, successful method for HD-DOT ^13,17,31,32^. Here, images were reconstructed by applying a regularized pseudoinverse of **A** to the simulated measurements **y**. Regularization used a spatially-variant Tikhonov (L2-norm) penalty. Thus, image reconstruction was equivalent to solving the optimization problem in the following equation to obtain an estimate 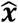 of the true underlying image **x**:

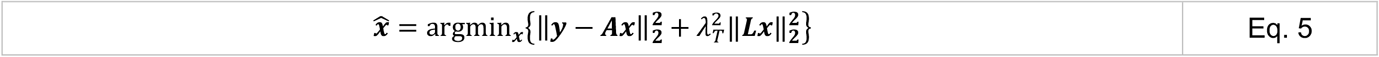

*λ*_*T*_ is the Tikhonov regularization parameter, a positive number that controls the strength of the regularization term’s influence. **L** is a diagonal matrix that incorporates the spatially variant regularization as follows, where *λ*_2_ is a positive number:

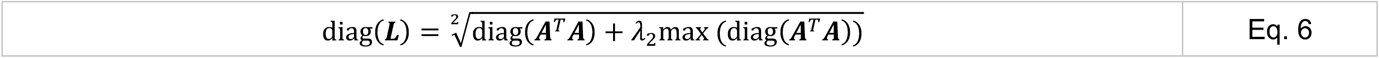

The reconstructed image was computed using a regularized Moore-Penrose pseudoinverse as follows, where 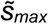 is the maximum singular value of **Ã**, where **Ã** = **AL**^-1^. The Tikhonov regularization parameter *λ*_*T*_ is set to 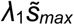 where *λ*_*1*_ is a positive number. This allows *λ*_1_ to be interpreted as a fraction of the maximum singular value of **Ã**.

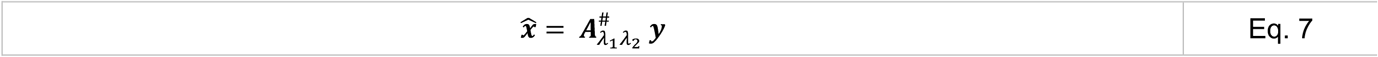

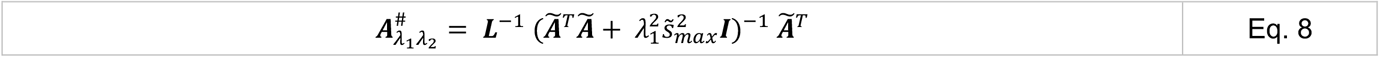

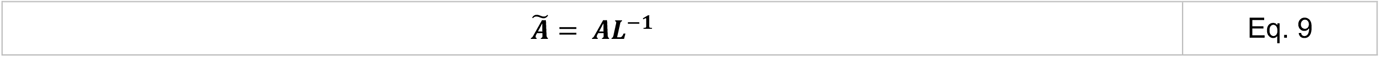

The parameter *λ*_1_ controls a key tradeoff between noise and spatial resolution in the reconstructed images. Smaller values of *λ*_1_ produce images with lower spatial resolution and more noise. This is due to the properties of **Ã** for DOT systems and the relationship between the regularized pseudoinverse and the singular value decomposition of **Ã**, as explained further in **Supplementary Information**.

The parameter *λ*_2_ controls spatially-variant regularization, which is necessary to precondition the matrix **A** into the matrix **Ã** to avoid biasing the activity in the reconstructed images toward the head surface. Further details on spatially-variant regularization are explained in previous work.^12,13,32^ Throughout this study, we used *λ*_2_ = 0.1 except for the simulations reported in Fig. 1-2. In those simulations, we used *λ*_2_ = 0.1 for the 13-mm and 10-mm-spaced grids, *λ*_2_ = 0.03 for the 6.5-mm-spaced grid, and *λ*_2_ = 0.01 for the 3.25-mm-spaced grid.

Because light intensity signals are generally low and noisy when measured from source-detector pairs farther than 40 mm apart in human imaging experiments, measurements with source-detector distances > 40 mm were removed from the image reconstruction stage. This was achieved by removing the corresponding rows of **Ã** and **y** before computing the regularized pseudoinverse and 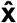 . We used the same distance cut-off when reconstructing images from in vivo DOT data to simplify comparative analyses.

#### M.1.3. Metrics of Image Quality

To evaluate the simulated performance of each array design, we calculated key measures of image quality including spatial resolution, localization error, effective resolution, and signal-to-noise ratio. These metrics were computed on the reconstructed images of simulated single-voxel perturbation at thousands of locations within the head for each array. The reconstructed image of each point target is termed the point-spread function (PSF) corresponding to that location.

Spatial resolution was quantified with the full-width at half-maximum (FWHM) of each PSF. To compute FWHM, first the maximum-value location (spatial peak) in each PSF was found. Second, voxels with values <50% of this maximum were discarded. Third, the remaining voxels were segmented into contiguous regions/clusters. The FWHM was computed as the maximum distance between voxels in the contiguous region *C* containing the maximum-value voxel and other voxels whose values were at least 50% of the maximum value. The localization error was quantified as the Euclidean distance between the perturbation location and the centroid of the reconstructed PSF volumetric image.

Effective resolution is a spatial uncertainty metric that combines spatial resolution and localization error ^11^. The effective resolution was computed as the diameter of the smallest sphere that is centered at the point perturbation location and that encloses the contiguous region of voxels with values ≥50% of the PSF’s maximum and containing the PSF’s maximum (i.e., region *C*).

The signal-to-noise ratio (SNR) was computed by identifying the voxels in region *C* of the PSF and dividing their average signal value by the average over voxels of their standard deviation across repetitions of the measurements. Further details on the SNR calculation are provided in **Supplementary Information**.

### M.2. Ultra-High-Density (UHD) DOT Imaging System Construction

#### M.2.1. Grid Configuration Overview

To leverage retinotopic paradigms for validation of image quality of occipital neural hemodynamic responses to visual stimuli, we constructed an “ultra-high-density” (UHD) DOT system with 6.5-mm nearest-neighbor spacing, 2 photon wavelengths, 126 source fibers, and 126 detector fibers placed in a regular grid array. The grid covered a 120 mm x 55 mm region over the back of the head, as in the simulation studies.

#### M.2.2. Two-Pass Encoding for Expanded Dynamic Range While Maintaining Minimal Crosstalk

To minimize crosstalk, we used a combination of frequency, temporal, and spatial encoding ^13^. For the frequency encoding, light sources at 685 nm and 830 nm were modulated at 13,228 Hz and 19,841 Hz, respectively. For temporal encoding, only one source position was illuminated (using laser diodes at both frequencies) at a time. To maintain a fast enough frame rate to oversample hemodynamic physiology, we also used spatial encoding such that identical temporal and frequency patterns were used simultaneously in sources that were separated by more than 9 cm to ensure no detector would be illuminated by both at a level above its noise floor. To circumvent detector saturation and achieve adequate dynamic range (10^6^-10^7^), we employed two-pass encoding into our illumination scheme. In the first pass through an encoding sequence, sources were illuminated for a longer amount of time using a 50% duty cycle than in the second pass that used a 1% duty cycle. The bright pass saturates nearby detectors but provides adequate light collection from farther-away detectors. The dim pass avoids saturating nearby detectors and is scaled to provide the equivalent light level as would have been seen with a 50% duty cycle. Further details on this scaling are provided in **Supplementary Information**.

#### M.2.3. Opto-electronics Infrastructure

Details of the UHD-DOT instrument are provided in **Supplementary Information**. To manage the increased inertia and weight due to the greater spatial density of optical fibers, we updated the design to fiber bundles with a slightly smaller cross section (from diameter 2.5 mm previously^13^ to 2.4 mm) and used PVC cladding instead of steel cladding. To increase source illumination efficiency and optical power, we updated from LEDs to laser diodes (LDs). We also upgraded analog-to-digital infrastructure to Focusrite Red A/D converters that use audio-over-ethernet communication and support 16 detectors per 1U of rack space. Encoded illumination patterns for the continuous wave (CW) HD-DOT system are delivered using two National Instruments PCIe-6537B high speed DIO cards (**Supplementary Fig. 3**). Each NI card has 32 output channels updated at a 10 MHz clock rate. To encode the 256 individual laser diode (LD) channels (128 source positions each with two independently encoded LDs) with 2x32 output channels, we temporally multiplexed the pattern data by a factor of four. These flashing patterns are demultiplexed using two custom “brain boxes” that each have four 32-channel output banks (A1, A2, B1, B2, C1, C2, and D1, D2 in **Supplementary Fig. 3**). The eight total output banks are each connected to custom source boxes using 40 pin ribbon cables. Each source box is equipped with power conditioning and 32 laser diode drivers (a 685-nm and an 830-nm LD per source position: Thorlabs HL6750MG and HL8338MG). The NIR light from the LDs is focused onto custom multimode source fibers (US FiberOptec 50-4256-1-REV1, 5.0 meters, single-core 400-μm cross section, NA = 0.48) using a custom optical SMA coupling. The optical fibers are bifurcated at the SMA end and terminate on the head with a 2.2 cm straight tip, 2.4 mm diameter ferule. The optical power measured at the end of the ferrule at the head is 23.5 ± 3.3 mW (mean ± standard deviation) for each 685-nm LD and 830-nm LD. Avalanche photo diodes (APDs, Hamamatsu, C12703-01 SPL) detect the light collected from the head via the detection optical fibers (US FiberOptec 50-4255-1-REV1, 5.0 meters, 2.4 mm diameter bundles of 30-micron fibers, NA = 0.66). After adjusting the gain of the low-noise current-to-voltage amplifier on the APD to approximately M = 90, the APDs have a noise equivalent power (NEP) = 19.1 ± 1.35 fW/√Hz, sensitivity = 4.9 ± 0.33 MV/W, and noise floor = 93.2 ± 3.3 nV in 1 Hz bandwidth. Custom built detector boxes (N = 22) regulate power and isolate signal for six APDs each. Each APD output is sampled at 96 kHz by a dedicated channel in a 24-bit, 12-channel A/D (Focusrite model A16R) with 2x10^-6^ crosstalk between channels. The remaining elements of the system were similar to those employed in previous work.^13^

#### M.2.4. Imaging Cap

The imaging cap and fiber management hardware were similar to a previous DOT system ^13^. However, the previous system’s nearest-neighbor optode spacing was 13 mm, which left more space for the rubber O-rings and 3D-printed mechanisms that held the optodes in place. To accommodate the shorter 6.5-mm inter-optode spacing of this system, the diameters of these components were reduced. This did not impact the rigidity of the cap or the components’ ability to hold the fibers in place and achieve adequate coupling to the scalp. Further details about the cap materials and fiber management are described in **Supplementary Information**.

### M.3. In-Vivo Human Imaging Experiments

#### M.3.1. Study Overview, Participants, and Stimuli

To evoke strong, repeatable brain responses in visual cortex, healthy adult human participants viewed flickering checkerboard wedge and ring stimuli at different polar and radial positions, respectively, during separate DOT and fMRI imaging sessions. The DOT data were collected from the UHD-DOT system. To evaluate the effect of optode grid density on in-vivo imaging performance, analyses were performed separately on data from the full UHD-DOT grid and with data from a subset of the optodes that formed a less-dense grid “a “high-density subset,” or “HDss,” of that grid). The nearest-neighbor spacing of this HDss grid was 14.5 mm.

Four healthy adult human participants completed this study. Participants were 20-33 years old, had no neurological conditions, and had normal or corrected-to-normal vision. All participants were right-handed females. All were recruited from Washington University and the St. Louis area. Informed consent was obtained from all participants. Research was approved by the Washington University Institutional Review Board (IRB) under IRB protocol #201707092 and was performed in accordance with this protocol.

The flickering checkerboard stimuli were similar to those employed in previous retinotopic mapping studies ^12,17,19,39^. The first stimulus type was a checkerboard wedge that subtended a 45° polar angle and 15° eccentricity in the visual field. This checkerboard wedge rotated clockwise or counterclockwise around the center of the screen, completed one full 360° revolution every 24 seconds and stepped 15° per second. The second type of stimulus was a logarithmic checkerboard ring that expanded outward or contracted inward. This was also presented periodically, cycling back to its initial position after reaching its final position. The ring swept through 12 positions covering the eccentricity range 1-15° in the visual field, completing 1 cycle every 24 seconds and spending 2 seconds at each position. All checkerboard stimuli were black and white and were overlaid on a 50% gray background. At a rate of 16 times per second, checkerboard color phase was inverted; i.e., black checks were switched to white checks and vice versa, yielding an 8-Hz “flicker” frequency. At all times, a white fixation cross was displayed on a gray background at the center of the screen, covering the central 2 degrees of the visual field. Participants were asked to keep their gaze fixated on this cross at all times.

Participants viewed these stimuli in runs, wherein only the fixation cross was displayed for 30 seconds, followed by the presentation of one type of checkerboard, moving in one direction, for at least 6 cycles (also called repetitions or blocks) with no breaks between cycles. Therefore, during each imaging session, participants viewed the clockwise wedges, counterclockwise wedges, expanding rings, and contracting rings each for a total of at least 144 seconds. During fMRI imaging sessions, subjects completed one run of each of those four run types with 6 cycles of the stimulus per run. During DOT imaging sessions, subjects completed three runs of each of the four run types with 13 cycles of the stimulus per run, with one exception: for two of the four run types (clockwise wedges and contracting rings), Subject 2 completed two runs with 13 stimulus cycles each, instead of three such runs. Between runs, participants remained in the scanner and were provided instructions and a brief description of stimuli for the next run.

#### M.3.2. DOT Data Preprocessing and Image Reconstruction

DOT data from each stimulus run were preprocessed in a similar manner as in prior studies ^13^. These preprocessing steps were designed to reduce signal components unrelated to brain activity. Briefly, these steps included conversion of the measured light level time trace into a relative change (log-ratio), rejection of noisy channels, regression of signals from superficial tissue (e.g., scalp), down-sampling to a 1-Hz frame rate, and band-pass filtering to 0.02-0.2 Hz. Images of optical absorption fluctuations *δμ*_*a*_ at each light wavelength were reconstructed from these preprocessed log-ratio time courses using Eq. 7, and these images were spectroscopically converted into oxygenated hemoglobin and deoxyhemoglobin concentration fluctuations. More details are provided in **Supplementary Information**.

#### M.3.3. MRI Data Acquisition and Preprocessing

To get a “ground-truth”/gold-standard measure of the evoked activity in visual cortex, we acquired fMRI data while subjects viewed the checkerboard stimuli. MRI scans were conducted on a Siemens Magnetom PRISMA Fit 3.0 T scanner, with an iPAT-compatible 20-channel head coil. Anatomical T1-weighted MPRAGE (echo time (TE) = 2.9 ms, repetition time (TR) = 2,500 ms, flip angle = 8°, 1 mm × 1 mm × 1 mm isotropic voxels, 256 × 256 × 176 voxel array/field of view) and T2-weighted (TE = 564 ms, TR = 3,200 ms, flip angle = 120°, 1 mm × 1 mm × 1 mm voxels, 256 × 256 × 176 voxel array/field of view) scans were collected for each participant at the beginning of each MRI scan session. Functional images were then collected using a series of asymmetric gradient spin-echo echo-planar (EPI) sequences (TE = 33 ms, TR = 1,230 ms, flip angle = 63°, 2.4 mm x 2.4 mm x 2.4 mm isotropic voxels, 90 × 90 × 64 voxel array/field of view, multi-band factor = 4) to measure the blood oxygenation level-dependent (BOLD) contrast.

MRI data were subjected to standard fMRI pre-processing as detailed previously ^13,40,41^, including correction of systematic slice-dependent time shifts, elimination of odd-even slice intensity differences to interleaved acquisition, rigid-body realignment for head motion across runs, normalization of signal intensity to a mode value of 1000, and segmentation of anatomical volumes using FreeSurfer ^33^. The anatomical images were segmented into scalp, skull, cerebrospinal fluid (CSF), gray matter, and white matter using FreeSurfer and NeuroDOT software, and this segmented image was employed to generate a UHD-DOT sensitivity matrix (**A** in Eq. 3) for each subject using NIRFAST and NeuroDOT software (section M.1.1).

The BOLD fMRI signal in each voxel was converted into percent BOLD change by subtracting and then dividing by the average BOLD signal in that voxel over time. Subsequently, temporal band-pass filtering (0.02-0.2 Hz) was applied, followed by nuisance regression with six rigid body values derived from head motion correction, white matter, CSF, and the mean whole-brain signal. Head motion was evaluated by the calculation of framewise displacement (FD) ^42-45^. Next, the percent BOLD change images from each human subject were affine-transformed into the subject’s native anatomical coordinate space defined by their T1-weighted MRI where the tissue segmentation, light propagation model, and UHD-DOT sensitivity matrix had been generated. This allowed all subsequent image processing and analyses to occur in each subject’s native space. Finally, these images were convolved with an isotropic Gaussian smoothing kernel with 10-mm FWHM, the same smoothing kernel that was applied to the DOT images after DOT image reconstruction.

#### M.3.4. Noise-Resolution Tradeoff Evaluation with Head Phantom and In-Vivo Data

To assess the noise vs. resolution tradeoff, we used three approaches. In the first approach, we sought to evaluate the effect of grid density on noise-resolution tradeoff in the absence of physiologic contributions to the noise. To accomplish this, a head phantom was imaged, and the noise was estimated from these data, while resolution was estimated from the FWHM of simulated PSF reconstructions, at different values of the regularization parameter *λ*_1_. The noise was calculated as the average, over voxels, of each voxel’s standard deviation over time points spaced 24 sec apart within the region under the optode grid footprint and extending 14 mm deep. The signal was calculated as the average, over voxels, of each voxel’s oxyhemoglobin signal in the contiguous region *C* containing the voxel with maximum oxyhemoglobin change and containing voxels with signal level ≥ 50% of that maximum in one of the block-averaged images of a human subject’s brain response to a checkerboard ring. The resolution was calculated as the FWHM of the simulated PSF image for a point target at the voxel with maximum signal level in the same block-averaged image of a human subject’s brain response to a checkerboard ring.

In the second approach, we sought to evaluate the effect of grid density on noise-resolution tradeoff in the presence of measured physiologic contributions to noise, but with controlled, simulated PSF images for resolution assessment. To accomplish this, noise in the images was calculated in the same way as for the head phantom, but using images from a human subject viewing the expanding checkerboard ring stimuli instead of images from the phantom. Signal and resolution were again quantified in the same ways from the simulated PSF reconstructions and were therefore identical to the signal and resolution values from the noise-resolution evaluation on the phantom.

In the third approach, we sought to evaluate the effect of grid density on noise-resolution tradeoff in fully in-vivo, human imaging. Here, resolution was quantified by the cube root of the volume (FVHM^1/3^) of the contiguous region *C* containing the maximum evoked oxyhemoglobin change in an image formed by dividing the average oxyhemoglobin change by its standard deviation over blocks (stimulus repetitions) within each voxel. SNR was obtained from the peak of that image’s transverse slice shown in **Fig. 5D**.

#### M.3.5. Evoked Response Feature Robustness Assessment

In the previous analysis, we focused on analyzing FVHM of a single activation. Most retinotopic responses are more complex than this. For example, in some places and times, the evoked activation appears to split into two distinct clusters. This provides another opportunity for evaluating resolution because a higher-resolution imaging system should see this splitting, whereas a lower-resolution system will blur the clusters together. We treated the fMRI data as a “ground-truth”/gold-standard measure of the activity evoked by the checkerboard stimuli and compared the block-averaged fMRI images with the block-averaged DOT images.

In addition to validating these complex DOT images by comparison to fMRI, we also validated them by assessing their repeatability across different subsets of the acquired data. To do this, we split our DOT data into three epochs of equal duration and computed block-averaged activation *t*-maps from these three subsets. Each *t*-map was computed by dividing the average oxyhemoglobin fluctuation signal by its standard error over blocks (stimulus repetitions) with each voxel (**Fig. 5A-B**).

#### M.3.6. Neural Decoding of Stimulus Position

A challenge with evaluating resolution using in vivo retinotopy paradigms is that the evoked activations are more spatially complex than simple points. Neural decoding tasks are also sensitive to the spatial information content (space-bandwidth product and SNR) of an imaging system. To supplement the noise-resolution analysis, we also evaluated the potential improvement from the higher-density grid in a neural decoding task. Therefore, we evaluated the performance of the two grids at discriminating the position/phase of the checkerboard wedge and ring stimuli using a simple brain activity decoding scheme ^21^.

In this decoding scheme, oxyhemoglobin images from two thirds of the repetitions of each stimulus were first block-averaged to form “templates,” forming one template image for each stimulus position. This was the training phase of the decoding algorithm. Then, in the testing phase, the spatial Pearson correlation was calculated between each template and the oxyhemoglobin image at each time point of the remaining one third of the repetitions of the stimulus. The scalp was excluded from the region of interest so that the spatial correlation was taken only over voxels in brain tissue and voxels in neighboring cerebrospinal fluid and skull tissues into which some of the brain activity image components may have been blurred or shifted. The decoding algorithm estimated the stimulus position at each such time point as the position whose template had the maximum correlation with that test time point. The accuracy of this decoding was quantified by the absolute difference between the true stimulus position and the decoder-guessed stimulus position. This process was repeated for twenty distinct training-testing splits: i.e., twenty ways of dividing the stimulus repetitions into training and testing sets containing two thirds and the other one third of the repetitions, respectively. The absolute difference between the true stimulus position and the decoder-guessed stimulus position (“decoding error”) was then averaged across test time points and across the twenty training-testing splits to obtain an overall metric of decoding accuracy.

#### M.3.7. Statistical Information

To evaluate the statistical significance of the UHD grid’s improved decoding accuracy, we performed a permutation test. In this test, we first computed the observed difference between the HDss and UHD grids’ decoding error at test time points spaced 13 seconds apart, and we averaged this observed difference across those time points and the twenty training-testing splits, an average over 480 time points. The spacing of 13 seconds was chosen to ensure that the values being averaged had negligible or no autocorrelation. Next, we estimated the null distribution of the mean HDss-UHD decoding error difference by randomly switching (permuting) the grid labels (UHD and HDss) at each time point with probability 0.5, then calculating the mean decoding error difference over time points in the permuted sample, and then repeating this for 10000 distinct permutations in total. The *p*-value was estimated as the fraction of null permutations in which the mean difference was greater than or equal to the observed mean difference. For cases where all 10000 permutations yielded a mean difference less than the observed mean difference, we simply concluded that *p* < 1/10000 (*p* < 10^-4^). Due to the Central Limit Theorem and our sample size (480 time points), the sampling distribution of the mean HDss-UHD decoding error difference was approximately Gaussian, so we estimated a 95% confidence interval on that mean using Student’s *t*-distribution with 480-1=479 degrees of freedom.^46^ We also computed effect size with Cohen’s *d* statistic.^47^ Detailed results of these statistical analyses are reported in **Supplementary Information** (section S10, table S1).

#### M.3.8. Ethical Approval for Human Experiments

Informed consent was obtained from all participants. Research was approved by the Washington University Institutional Review Board (IRB) under IRB protocol #201707092 and was performed in accordance with this protocol.

## ADDITIONAL INFORMATION

### Data Availability

Data collected and analyzed in this study are available from the corresponding authors upon reasonable request.

## AUTHOR CONTRIBUTIONS

Z.E.M., J.W.T., E.J.R., M.A.C., E.M.M., A.T.E., and J.P.C. conceived and designed the experiments. Z.E.M., J.W.T., E.J.R., K.T., S.M.R., A.M.S., M.L.S., T.M.B-Y., K.M.B., and A.T.E. performed the experiments. Z.E.M., J.W.T., E.J.R., K.T., A.T.E., and J.P.C. analyzed the data. Z.E.M., J.W.T., E.J.R., K.T., S.M.R., T.M.B-Y., K.M.B., A.T.E., and J.P.C. contributed materials/analysis tools. Z.E.M., J.W.T., E.J.R., K.T., A.T.E., and J.P.C. wrote the paper.

## ACKNOWLEDGMENTS

The authors thank Valinda K. Hood for her assistance with MRI scanner operation. In addition, we thank Patrick Mineault and Michael A. Choma for helpful conversations about this work. We also gratefully acknowledge funding from the following sources: National Institutes of Health grants U01EB027005, R01NS090874, K01MH103594, and F31NS110261, a Meta Sponsored Academic Research Agreement, and Washington University’s Cognitive, Computational, and Systems Neuroscience Fellowships.

