## Supplementary Information for "Ultra-high density imaging arrays for diffuse optical tomography of human brain improve resolution, signal-to-noise, and information decoding"

### S1. Noise Model for Simulation Studies

To simulate the noise **n** in the imaging system forward model (Eq. 4), we built a simplified noise model from in-vivo imaging data from an existing HD-DOT system with 13-mm inter-optode spacing [1]. Our goal was to model **n** as a random vector from a Gaussian probability distribution whose variances depended on the source-detector distances and simulated baseline light levels in each measurement channel so that **n** could be simulated for any arbitrary optode grid, as required for our simulation studies.

This was challenging because many sources of noise contribute to the light level time trace measured by each channel (source-detector pair at each light wavelength). Noise sources include but are not limited to source modulation noise, electronic detector noise, optode-scalp coupling changes due to head motion, and physiological background signals, such as cardiac pulse, respiration, and vasomotion, that are not part of the stimulus- and brain activity-evoked hemodynamics. Modeling all these factors mechanistically from physics and first principles would have been impractical, inaccurate, and beyond the scope of this study.

Therefore, we developed a data-driven, top-down model that related the source-detector distance $R_{SD,i}$ and simulated baseline light level $G_{SD,i}$ of a measurement channel ($i$) to the standard deviation (SD) of the preprocessed log-ratio time trace $y_i$ in that channel. The measurements were taken from a resting-state imaging session where the subject only viewed a fixation cross and performed no other task, so that this SD would amalgamate all sources of noise without including stimulus-related signal components, and therefore the SD of $y_i$ would equal the SD of the noise component $n_i$. This SD was also equivalent to the root-mean-square amplitude (RMS) of $y_i$ over time because the mean of each log-ratio time trace was subtracted during preprocessing. The noise in different channels was treated as uncorrelated because the amount of available data was insufficient to reliably estimate the full noise covariance matrix.

To build the noise model, we first fitted a log-linear relationship between the simulated baseline light level $G_{SD}$ and the source-detector distance $R_{SD}$ on a sphere with a UHD-DOT grid (Eq. S1, Supplementary Fig. 1A). The slope and intercept of this relationship are dictated by the tissue optical properties and the optical diffusion equation, not the density of the optode grid.[2-5] We then calculated a simple ratio relationship between $G_{SD}$ and the measured baseline light level $\Phi_0$ from an HD-DOT system (Eq. S2, Supplementary Fig. 1B). This ratio $\rho$ was calculated as the exponentiated mean measured baseline log light level divided by the exponentiated mean log

$G_{SD}$ value for source-detector distances 23-29 mm (Eq. S3). Next, we fitted a power-law relationship between the RMS of $y_i$ and the baseline light level $\Phi_0$ in the $i$-th channel (Eq. S3, Supplementary Fig. 1C). Finally, we combined these relationships to obtain a formula for estimating the RMS of $y_i$ from any $R_{SD}$ (Eq. S4). This also served as the formula for the SD of $n_i$ because for this dataset, the RMS of $y_i$ and the SD of $n_i$ were equal. We then generated each element $n_i$ of each simulated noise vector **n** as needed from a univariate Gaussian distribution with mean 0 and SD given by Eq. S5.

| | |
|---|---|
| $G_{SD}(R_{SD}) = 0.04e^{-0.26R_{SD}}$ | Eq. S1 |
| $\Phi_0(R_{SD}) = \rho G_{SD}(R_{SD})$ | Eq. S2 |
| $\rho = \frac{\exp\left[\mathrm{mean}_{23\mathrm{mm}<R_{SD}<29\mathrm{mm}} \ln\left(\Phi_0(R_{SD})\right)\right]}{\exp\left[\mathrm{mean}_{23\mathrm{mm}<R_{SD}<29\mathrm{mm}} \ln\left(G_{SD}(R_{SD})\right)\right]} = 3300$ | Eq. S3 |
| $RMS(y_i) = (7.7 \times 10^{-4})\Phi_0^{-0.14}$ | Eq. S4 |
| $RMS(n_i) = RMS(y_i) = (7.7 \times 10^{-4})(3300 \times 0.04e^{-0.26R_{SD}})^{-0.14}$ | Eq. S5 |

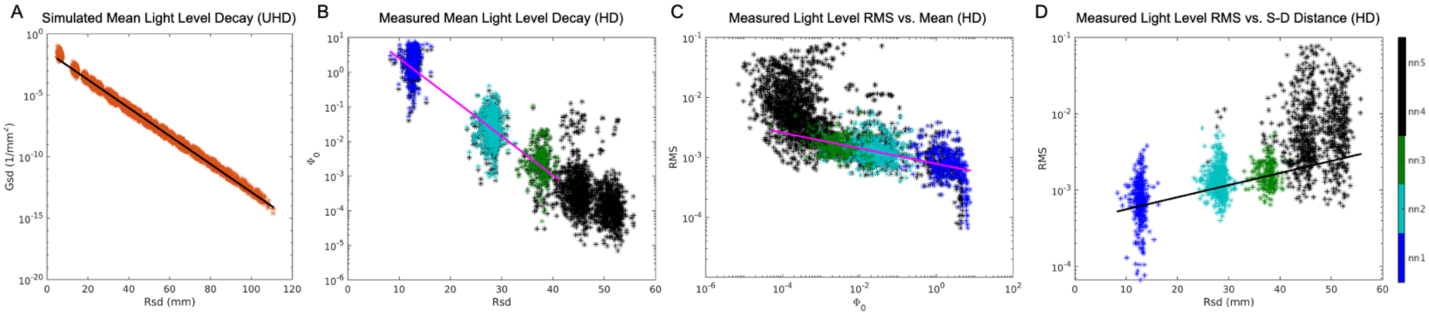


**Supplementary Fig. 1:** Developing a noise model by fitting relationships between source-detector distance, simulated baseline light level, measured baseline light level, and RMS of log-ratio measurement time traces. (A) Simulated baseline light level follows a log-linear decay with source-detector distance in the regime of distances present in the UHD-DOT grid. (B) Measured baseline light level also follows a log-linear decay with source-detector distance in the regime of distances present in the HD-DOT grid. (C) RMS of each measurement channel's log-ratio time trace is related to the channel's measured baseline light level via a power law. (D) RMS of each measurement channel's log-ratio vs. source-detector distance is well-fitted by the resulting noise model (Eq. S5). Black data points came from channels with source-detector distances > 40 mm or RMS > 0.075 and were excluded from the fitting procedure because these measurements are ignored in practice during image reconstruction due to their low signal-to-noise ratio.

### S2. Calculation of Signal-to-Noise Ratio (SNR) in Simulation Studies

For a given point-spread function (PSF), let region $C$ be the contiguous region containing the maximum-value voxel and other voxels whose values were at least 50% of the maximum value. The signal-to-noise ratio (SNR) was computed by identifying the voxels in region $C$ and dividing their average signal value by the average over voxels of their standard deviation across repetitions of the measurements. This is shown in the following equation, where $k$ is a voxel index, $p$ is an index denoting one repetition/instance of the PSF image (corresponding to one instance of noise **n**), $N_C$ is the number of voxels in region $C$, and $N_P$ is the number of repetitions of the measurements:

| | |
|---|---|
| $SNR = \frac{mean_{k\in C} mean_p \hat{x}_{k,p}}{mean_{k\in C}\sigma_k} = \frac{\frac{1}{N_C}\sum_{k\in C}\frac{1}{N_P}\sum_p \hat{x}_{k,p}}{\frac{1}{N_C}\sum_{k\in C}\sqrt{\frac{1}{N_P-1}\sum_p\left(\hat{x}_{k,p} - mean_p\left(\hat{x}_{k,p}\right)\right)^2}}$ | Eq. S6 |

### S3. Noise-Resolution Tradeoff Controlled by Tikhonov Regularization Parameter

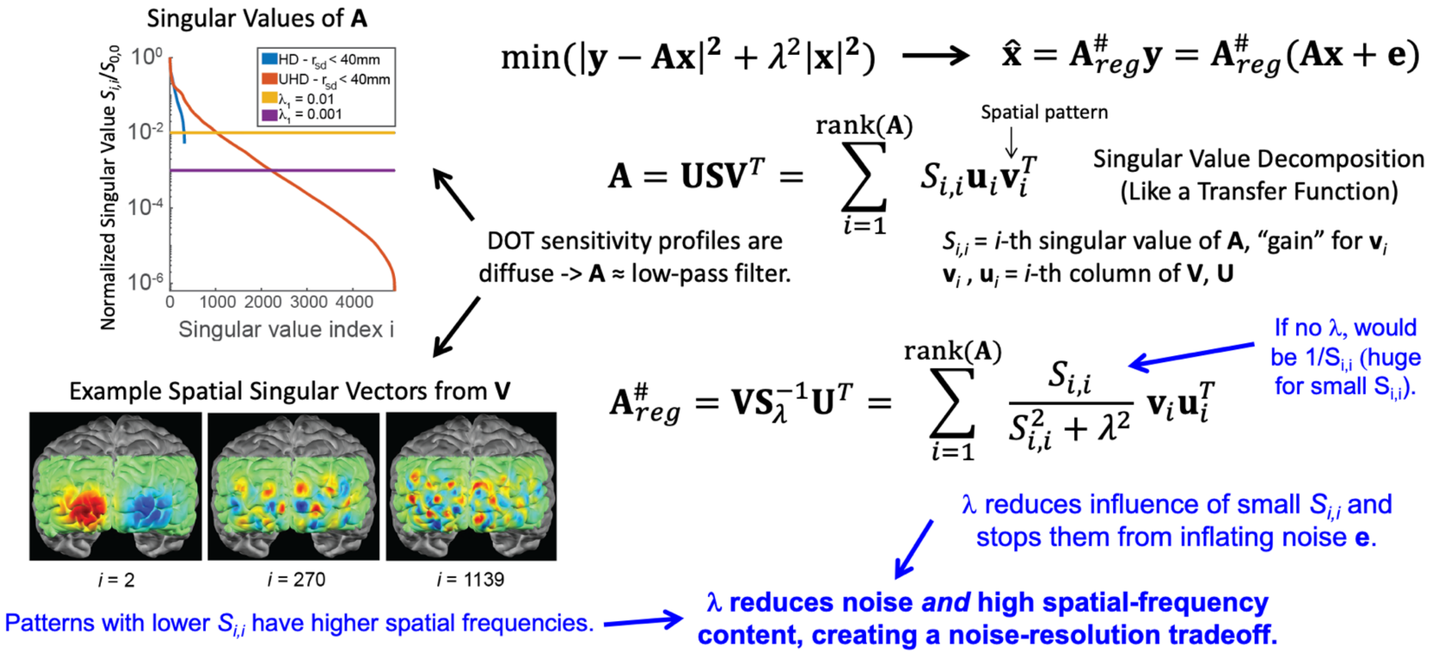


**Supplementary Fig. 2:** Tikhonov regularization parameter, proportional to $\lambda_1$, creates a tradeoff between image resolution and noise amplitude due to the singular values and vectors of the sensitivity matrix **A**. Singular vectors with higher singular values contain lower-spatial-frequency patterns because DOT measurements are more sensitive to spatially broader brain activations, due to the diffuse sensitivity profiles and frequent scattering of light in the tissue.

### S4. Two-Pass Encoding and Associated Scaling

To circumvent detector saturation and achieve adequate dynamic range ($10^6$-$10^7$), we employed two-pass encoding into our illumination scheme. In the first pass through an encoding sequence, sources were illuminated for a longer amount of time using a 50% duty cycle than in the second pass that used a 1% duty cycle. The bright pass saturates nearby detectors but provides adequate light collection from farther-away detectors. The dim pass avoids saturating nearby detectors and is scaled to provide the equivalent light level as would have been seen with a 50% duty cycle according to Eq. S7, where the duty cycles $DC_{\text{bright}}$ = 0.5 and $DC_{\text{dim}}$ = 0.01 and where $\Phi_{\text{bright}}$ and $\Phi_{\text{dim}}$ are the source intensities in the bright and dim pass, respectively.

| | |
|---|---|
| $$\Phi_{\text{bright}} = \Phi_{\text{dim}} \frac{\sin(\pi DC_{\text{bright}})}{\sin(\pi DC_{dim})}$$ | Eq. S7 |

All measurement pairs that clipped during the bright pass were replaced by the scaled data from the dim pass according to Eq. S7. By using the measurements from the bright pass for long source-detector separations and the measurements from the dim pass for short source-detector separations, a reliable overall dynamic range of ~$10^7$ was achieved (**Fig. 3d**). With the 2-pass encoding scheme and 91 temporal encoding steps, this encoding strategy sampled the field of view at a frame rate 7.33 Hz.

### S5. Opto-Electronics Infrastructure Schematic

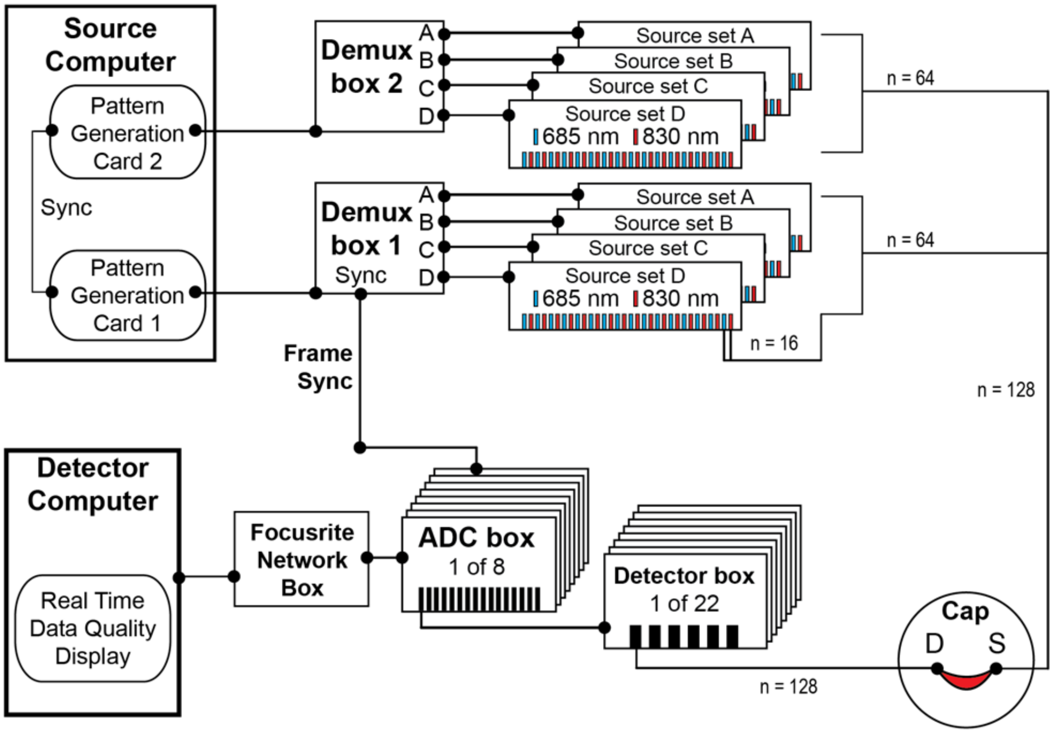


**Supplementary Fig. 3:** Connectivity of opto-electronic components for controlling and synchronizing light sources and detectors, recording measured light levels, and displaying real-time measures of data quality.

### S6. Imaging Cap Design and Materials

The imaging cap hardware and fiber suspension mechanism are similar to a previous DOT system [1]. The rigid backbone of the imaging cap is a layer of aquaplast that has been shaped to approximate the curvature of the head surface. The aquaplast (black) provides structural integrity with flexibility that is able to conform to a wide variety of head shapes and sizes during the cap fit. Soft foam (dark grey) lines the inside of the cap for greater comfort. The optic fiber tip is guided by a 'top-hat' style spacer (green) and can translate (arrows on light grey optode) perpendicularly to the head surface (textured yellow). The spacer is held in place with a rubber O-ring on the outside surface of the cap (orange). Foam O-rings (brown and red), snug-fit onto the tip of the optode, maintain a 3 mm penetration length of the fiber tip (for combing through hair), and provide a springy tension that is leveraged for a uniform cap fit over the curved and irregular shape of the subject's head. The cap is attached to the subject with hook-and-loop straps (blue): two positioned on the forehead and two on the top of the head (to conform the curvature of the cap to a wide variety of head morphologies). The top-hat spacers along with the locally-rigid aquaplast minimize torque due to forces parallel to the head surface.

The fiber management around the head provides lateral force (perpendicular to the surface of the head) on each optode that couples the fiber tip against the scalp of the subject. The fibers are supported from above in a double-halo design that uniformly counter-balances the weight of the fibers around the head, and fully removes strain of fiber weight from the subject. Supporting the weight of the fibers, the double-halo, and the cap well above the subject (supports are attached > 1 m above the cap), allows the cap to move as a pendulum free to swing in two dimensions. This freedom minimizes coupling perturbations at the fiber scalp coupling interface caused by subject motion.

### S7. DOT Data Preprocessing and Image Reconstruction

DOT data from each stimulus run were preprocessed similarly to prior studies to reduce signal components unrelated to brain activity [1]. The raw light level time trace $\Phi(t)$ in each measurement channel was converted into a fractional change by log-ratio transformation: i.e., the light levels were divided by their mean value over time, and the natural logarithm of this result was computed to obtain the optical density $y(t)$. Channels whose log-ratio trace had standard deviation > 0.075 over time (7.5% of mean light level) were labeled as noisy and were excluded from further processing. For each channel, the linear trend over time was removed, followed by band-pass filtering (pass band of 0.02-1.0 Hz) to remove long-term drifts and high-frequency noise unrelated to the checkerboard stimuli. Next, an average time trace over all measurement channels with source-detector distance < 15 mm (sampling predominantly scalp and skull) was constructed as an estimate of systemic and superficial signals. This estimate of the superficial signal time course was then linearly regressed out of every measurement channel time trace. Then a final low-pass filter with cutoff 0.2 Hz was applied to each channel to remove remaining non-neural signal components unrelated to the checkerboard stimuli, such as variance due to cardiac cycles and respiration, ultimately yielding signals with frequency content in the 0.02-0.2 Hz band. The data were then temporally down-sampled to a 1-Hz frame rate.

Images of optical absorption fluctuations $\delta\mu_a$ at each light wavelength were reconstructed from these preprocessed log-ratio time courses using Eq. 7 and a sensitivity matrix **A** built from each participant's anatomical MRI images using NIRFAST, FreeSurfer, and NeuroDOT software as previously described [1]. To reduce image noise, each volume was then spatially smoothed with an isotropic Gaussian kernel with 10-mm FWHM. Finally, the optical absorption fluctuations at the two light wavelengths were spectroscopically converted into oxygenated hemoglobin and deoxyhemoglobin concentration fluctuations $\delta Hb_h$ using Eq. S8 below, where $\boldsymbol{E}$ is the extinction coefficient matrix, $w$ is an index denoting optical wavelength, and $h$ is an index denoting oxy- or deoxyhemoglobin:

| | |
|---|---|
| $\delta Hb_h = \sum_{w=1}^{2} (\boldsymbol{E}^{-1})_{h,w} (\delta\mu_a)_w \qquad \boldsymbol{\delta H} = \boldsymbol{E}^{-1}\boldsymbol{\delta\mu_a}$ | Eq. S8 |

**S8. Phase-Encoded Retinotopic Mapping**

To further validate the UHD-DOT system, we estimated retinotopic maps from our DOT images. These retinotopic maps indicate which part of the stimulus visual field most strongly activates the tissue in each voxel of the brain. Retinotopic maps are useful for validation because they have an expected organization based previous brain imaging and electrophysiological studies. For example, voxels in the left brain hemisphere should be most sensitive to stimuli in the right visual hemifield and vice versa, and roughly speaking, voxels more superior and anterior within a visual area should be more sensitive to stimuli farther from the center of the visual field.[6-8]

To estimate retinotopic maps from our UHD-DOT images, we exploited the periodicity of our stimuli. Because our stimuli were periodic in time, stimulus phase within the repetition block corresponds to stimulus position, so the phase of a voxel's activation at the stimulus repetition frequency indicates which stimulus position most strongly activates that voxel's tissue. Therefore, we Fourier-transformed each voxel's time trace to compute its phase $\psi_{fw}$ at the stimulus repetition frequency $f_{st}$ ($24^{-1}$ $\text{sec}^{-1}$ = 0.0417 Hz) over a run with many repetitions of the stimulus moving forward through its cycle. Due to the time-delay between the stimulus-evoked electrical activity and the resulting blood oxygen fluctuation in the brain (the hemodynamic response function),[9-11] the DOT voxel time traces' phases were inherently delayed relative to the stimulus phase/position. To estimate this phase delay $\Delta\psi_{HDR}$ for each voxel and accordingly shift the voxel's phase to match the stimulus position, we also computed each voxel time trace's Fourier phase $\psi_{bw}$ at the stimulus repetition frequency from a run where the stimulus moved backward through its cycle. Defining $\psi'$ as the true stimulus phase to which a voxel is most sensitive, we employed the simple model in Eq. S9 and Eq. S10 and solved for $\psi'$ and $\Delta\psi_{HDR}$ independently for each voxel.

| | |
|---|---|
| $\psi_{fw} = \psi' + \Delta\psi_{HRF}$ | Eq. S9 |

| | |
|---|---|
| $\psi_{bw} = -\psi' + \Delta\psi_{HRF}$ | Eq. S10 |

We repeated these calculations over three pairs of forward and backward runs through the checkerboard wedge stimulus to produce the retinotopic maps of $\psi'$ in Supplementary Fig. 4. Because $\psi'$ is attributable to the stimulus position only in voxels that adequately respond to the stimulus, we computed values of $\psi'$ only for voxels whose signal power at the stimulus repetition frequency was at least 2.5% times the maximum power at that frequency on the brain surface within the imaging system's field of view. Broadly, these maps were repeatable when estimated from different runs (repetition groups), and these maps displayed the expected retinotopic organization patterns from previous studies of visual cortex.[6-8,12] In addition, the maps from the UHD-DOT system

(6.5-mm-spaced grid) appear sharper than the maps from the HD (13-mm-spaced) subset of that grid, consistent with the expected higher resolution of UHD-DOT. However, the maps from subject 4 were noisy and inconsistent between different subsets of the data. This is likely due to the low raw data quality obtained from subject 4, potentially due to poor optode-scalp coupling (poor cap fit) or thick skull and scalp; see section S9 for further discussion of raw data quality in each subject.

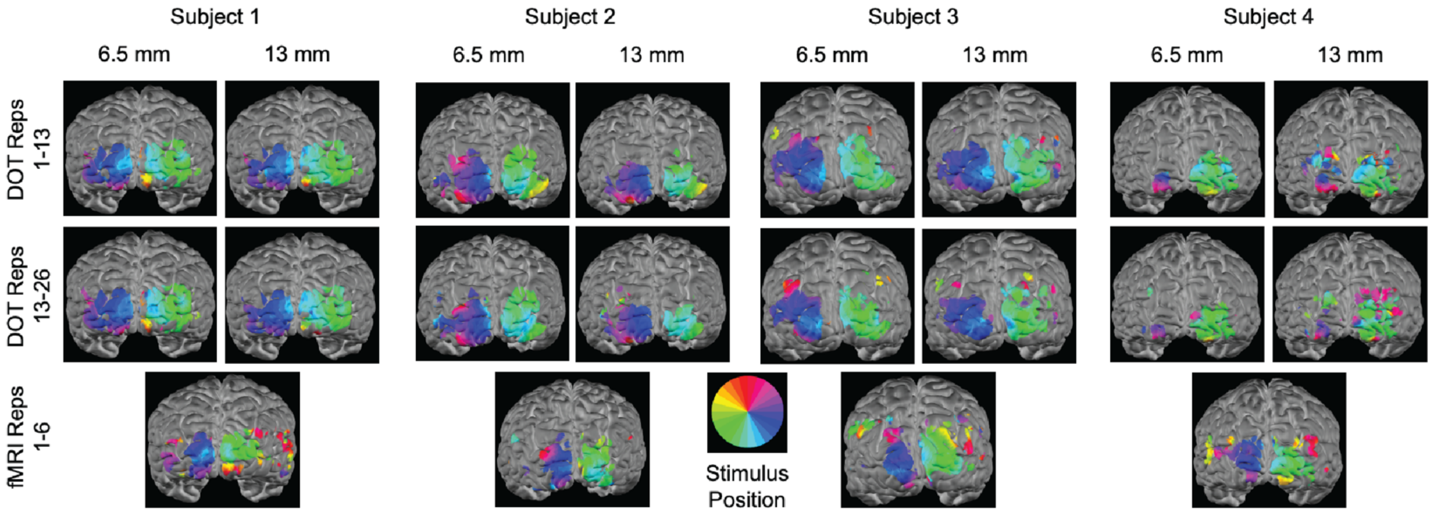


**Supplementary Fig. 4:** Phase-encoded retinotopic maps indicating which stimulus polar angle position in the visual field most strongly activates each brain voxel. These maps were repeatable across different subsets (repetition groups) of the data and were consistent with previously observed retinotopic organization patterns in these brain areas. UHD-DOT (6.5-mm-spaced grid) produced retinotopic maps that were less blurry and contained more of the qualitative features of the fMRI retinotopic maps than the HD (13-mm-spaced) subset of the UHD grid.

### S9. Cardiac Pulse Signal-to-Noise Ratio

A key metric of data quality is cardiac pulse signal-to-noise ratio because in a human subject, the cardiac pulse should be strongly visible in the hemodynamic signals that DOT seeks to measure. If the cardiac pulse is not strongly present in measured data, then the raw data quality is poor and unlikely to reliably reflect neural activity even after filtering, likely due to weak optode-to-scalp coupling/contact (poor cap fit), subject head motion, and/or thick skull and scalp tissue. In contrast, in a high-quality dataset, the cardiac pulse will produce a peak in the time traces' power spectra in the 0.5-2.0 Hz band, and the area under this peak (pulse signal power $P_{CS}$) will far-exceed the median area under the power spectrum in neighboring bands of width equal to the pulse peak width ("noise" or background signal power $P_{CBG}$ with respect to the cardiac pulse). We define a time trace's cardiac pulse SNR ($SNR_{CP}$), in dB, as 10x the log of the ratio of these two powers (Eq. S11).

$$SNR_{CP}\,(dB) = (10\,dB)\cdot\log_{10}\frac{P_{CS}}{P_{CBG}} \qquad \text{Eq. S11}$$

Supplementary Fig. 5 shows the average cardiac pulse SNR in each UHD-DOT optode for each subject in different runs (stimulus repetition groups). The average cardiac pulse SNR in each optode was obtained by averaging the cardiac pulse SNR in each measurement channel containing that optode and having source-detector distance 21-35 mm. Measurement channels labeled as extremely noisy during preprocessing (i.e., measurement channels with temporal SD > 0.075 in their log-ratio time traces) were excluded from the cardiac pulse SNR calculations.

In subjects 1-3, nearly all optodes had average cardiac pulse SNR ≥ 10 dB (≥1 order of magnitude), indicating reasonable data quality. These SNR measures were also similar in these two subsets of the data. In contrast, subject 4 had cardiac pulse SNR < 6 dB in all optodes, with about 25% of optodes having cardiac pulse SNR within roundoff error of 0 (gray optodes), indicating poor data quality. This is likely why the data from subject 4 produced erratic retinotopic phase maps and high stimulus position decoding error.

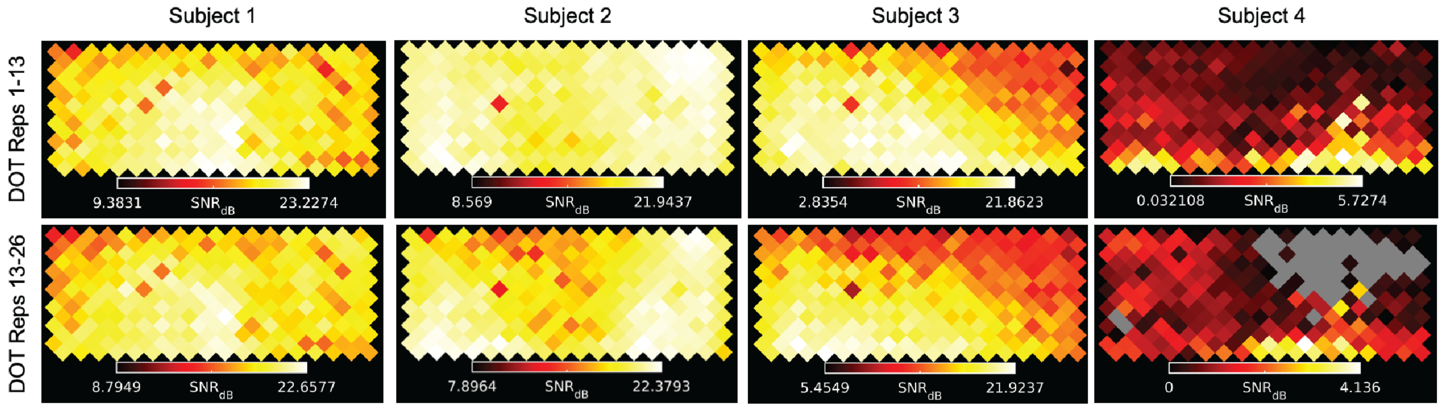


**Supplementary Fig. 5:** Average cardiac pulse SNR in each optode of the UHD-DOT cap in each subject, calculated from two different runs (stimulus repetition groups) of the clockwise-rotating wedge stimulus. In subjects 1-3, the cardiac pulse was ≥10x stronger than background signals in adjacent frequency bands, indicating reasonable data quality. However, in subject 4, the cardiac pulse was <3x stronger than background signals in nearly all optodes, with 25% of optodes having undetectable cardiac pulse, indicating poor data quality and explaining the erratic retinotopic maps and high stimulus decoding error seen in subject 4.

### S10. Detailed Illustration of Stimulus Position Decoding and Statistical Testing

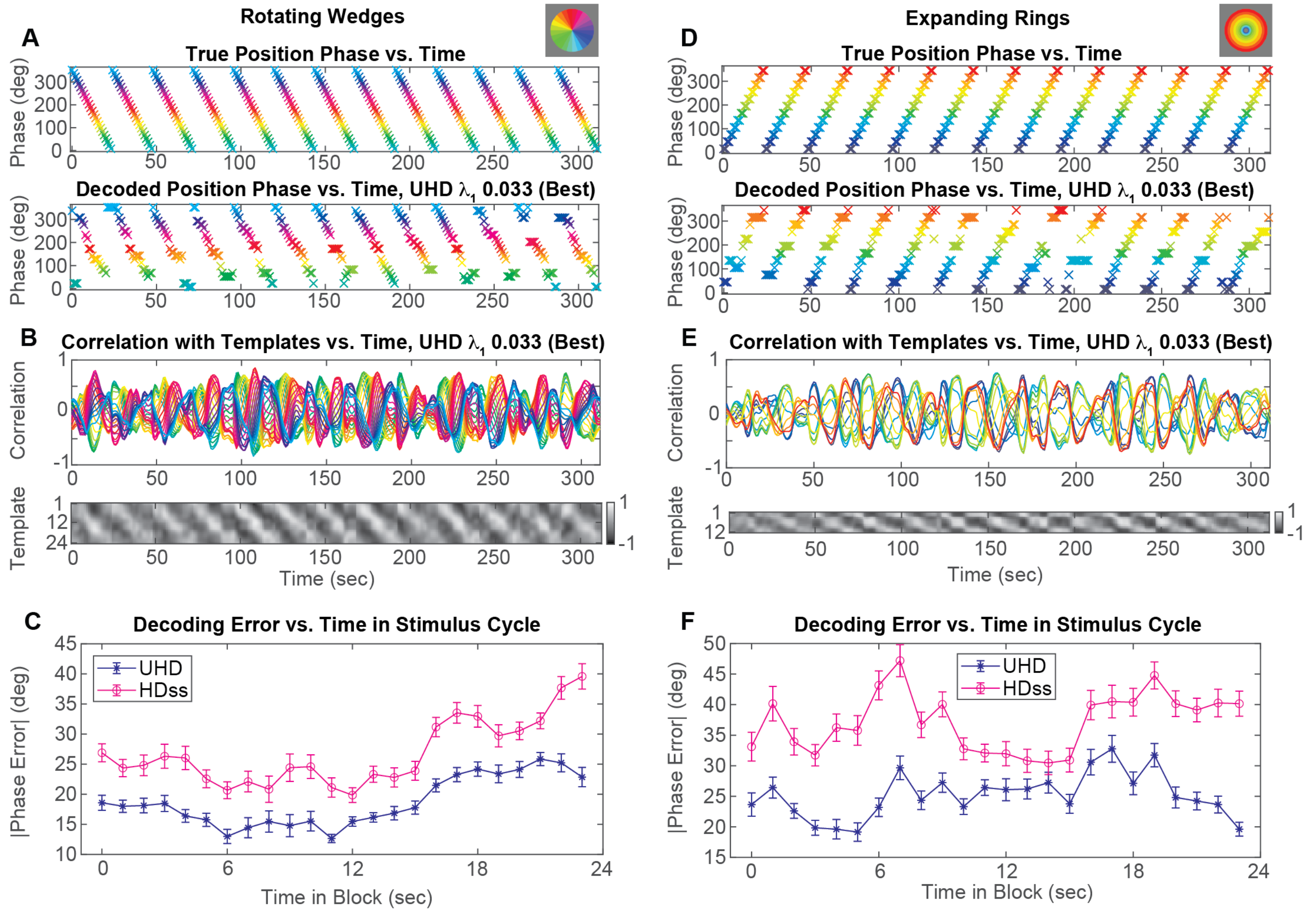


**Supplementary Fig. 6:** Stimulus decoding via template-based, maximum-correlation classification, illustrated for subject 3. (A) True and decoded wedge position vs. time within a test run, for the UHD grid at the $\lambda_1$ value that yielded lowest decoding error for that grid in this subject. (B) Spatial Pearson correlation between each template from the training phase and the oxyhemoglobin image at each time point in the testing phase. (C) Absolute error in decoded wedge position, averaged over repetitions/blocks of the wedge presentation. Error bars denote standard error of the mean for this average absolute error over blocks. The horizontal axis is the time within a block (block duration = 24 seconds). For all stimulus positions (times within each block), the UHD grid more accurately decoded the stimulus position than the HD-subset grid. (D-F) Analogous to panels A-C but for the expanding ring stimuli.

| Subject | Stimulus | Mean HDss-UHD Decoding Error (deg) | 95% Confidence Interval (deg) | Degrees of Freedom | Permutation Test p-Value | Effect Size (Cohen's d) |
|---|---|---|---|---|---|---|
| 1 | Wedges | 2.59 | [1.64, 3.55] | 479 | $<10^{-4}$ | 0.243 |
| | Rings | 2.69 | [1.61, 3.77] | 479 | $<10^{-4}$ | 0.223 |
| 2 | Wedges | 6.62 | [4.66, 8.59] | 479 | $<10^{-4}$ | 0.303 |
| | Rings | 4.06 | [2.21, 5.92] | 479 | $<10^{-4}$ | 0.196 |
| 3 | Wedges | 6.56 | [4.70, 8.43] | 479 | $<10^{-4}$ | 0.316 |
| | Rings | 9.00 | [6.49, 11.5] | 479 | $<10^{-4}$ | 0.322 |
| 4 | Wedges | 17.5 | [13.7, 21.3] | 479 | $<10^{-4}$ | 0.418 |
| | Rings | 16.2 | [12.5, 19.9] | 479 | $<10^{-4}$ | 0.396 |

**Table S1:** Statistical testing of stimulus position decoding error differences between UHD and HDss grids indicates clear, robust improvements in decoding accuracy from the UHD grid. These statistics were calculated as described in Methods section M.3.6.

### S11. Illumination Encoding Dwell Time Considerations in Comparing Grids

In several of our analyses of experimental data, we compared images from the full 6.5-mm-spaced grid to images from a subset of those channels at 13-mm spacing. While these comparative assessments of 6.5-mm and 13-mm arrays have the advantage of being simultaneous, a separate 13-mm system would have fewer time encode steps and longer dwell times during illumination. However, while this would likely improve the static phantom data, which is dominated by electro-optic noise, because the in-vivo variance 5x larger than phantom variance (**Fig. 4**), it is highly unlikely that the in-vivo performance of the 13-mm grid measurements would improve with the longer dwell times.
